# Ancestral protein reconstruction reveals evolutionary events governing variation in Dicer helicase function

**DOI:** 10.1101/2022.12.30.522297

**Authors:** Adedeji M. Aderounmu, P. Joseph Aruscavage, Bryan Kolaczkowski, Brenda L. Bass

## Abstract

Antiviral defense in ecdysozoan invertebrates requires Dicer with a helicase domain capable of ATP hydrolysis. But despite well-conserved ATPase motifs, human Dicer is incapable of ATP hydrolysis, consistent with a muted role in antiviral defense. To investigate this enigma, we used ancestral protein reconstruction to resurrect Dicer’s helicase in animals and trace the evolutionary trajectory of ATP hydrolysis. Biochemical assays indicated ancient Dicer possessed ATPase function, that like extant invertebrate Dicers, is stimulated by dsRNA. Analyses revealed that dsRNA stimulates ATPase activity by increasing ATP affinity, reflected in Michaelis constants. Deuterostome Dicer-1 ancestor, while exhibiting lower dsRNA affinity, retained ATPase activity; importantly, ATPase activity was undetectable in the vertebrate Dicer-1 ancestor, which had even lower dsRNA affinity. Reverting residues in the ATP hydrolysis pocket was insufficient to rescue hydrolysis, but including additional substitutions distant from the ATPase pocket rescued vertebrate Dicer-1’s ATPase function. Our work suggests Dicer lost ATPase function in the vertebrate ancestor due to loss of ATP affinity, involving motifs distant from the active site, important for coupling dsRNA binding to the active conformation. RLRs important for interferon signaling, and their competition with Dicer for viral dsRNAs, possibly provided incentive to jettison an active helicase in vertebrate Dicer.

## Introduction

Dicer is a multidomain endoribonuclease that is conserved in most eukaryotes^1–6^. Some organisms encode only a single Dicer^6–13^, while others encode multiple Dicers^3,4,14,15^, with different versions specialized for pre-microRNA (pre-miRNA) processing or endogenous/viral double-stranded (dsRNA) processing^14–23^. Dicer contains an intramolecular dimer of two RNaseIII domains, the catalytic center that cleaves dsRNA. It also contains a platform/PAZ domain, an N-terminal helicase domain, a C-terminal dsRNA-binding motif (dsRBM) and a domain of unknown function (DUF283) with a degenerate dsRBM fold^6,17,24^ (Figure 1A). These domains mediate recognition, binding, and discrimination of different dsRNAs, ensuring that optimal Dicer substrates are presented to the catalytic center for cleavage. The size of the small RNA product, either miRNA or small interfering RNA (siRNA), is defined by the distance between the platform/PAZ domain, which binds the ends of dsRNAs, and the RNaseIII domains^16,25,26^. Like the platform/PAZ domain, Dicer’s helicase domain is capable of binding dsRNA termini, and in some organisms, the C-terminal dsRBM contributes to substrate binding and cleavage^16,27,28^. Because both domains bind dsRNA termini, there is potential for the platform/PAZ and the helicase to compete for dsRNA substrates. To resolve this conflict, some extant metazoan Dicers have evolved substrate preferences where the platform/PAZ domain is specialized for binding the 2-nucleotide (nt) 3’ overhang (3’ovr) of a pre-miRNA, while the helicase prefers dsRNA with blunt (BLT) termini^29,30^.

**Figure 1.**
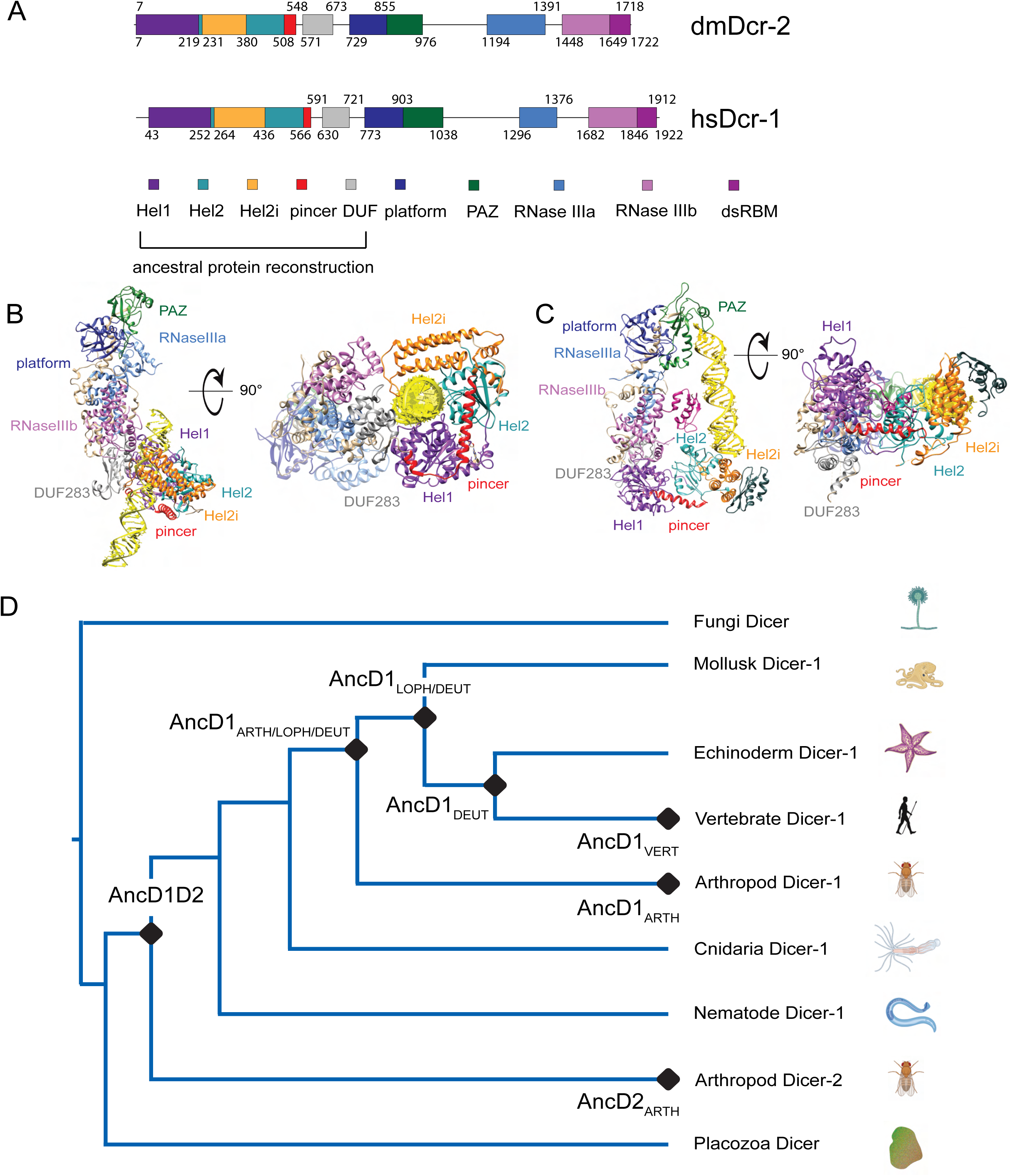
Phylogenetic analysis of Helicase domains and DUF283 of metazoan Dicer proteins. **A.** Domain organization of *Drosophila melanogaster* Dicer-2 and *Homo sapiens* Dicer, with colored rectangles showing conserved domain boundaries indicated by amino acid number. Domain boundaries were defined by information from NCBI Conserved Domains Database (CDD), available crystal and cryo-EM structures, and structure-based alignments from previous studies^24,28,76^. **B.** Structure of dmDcr2 bound to dsRNA (yellow) at the helicase domain. Left: front view. Right: bottom-up view. (PDB: 7W0C). **C.** Structure of hsDcr bound to dsRNA (yellow) at the platform/ PAZ domain. Left: front view. Right: bottom-up view. (PDB: 5ZAL). **D.** Summarized maximum likelihood phylogenetic tree constructed from metazoan Dicer helicase domains and DUF283. Nodes of interest are indicated with black rounded rhombi. ARTH: Arthropod, LOPH: Lophotrochozoa, DEUT: Deuterostome, VERT: Vertebrate.

The role of the helicase domain varies between different Dicers. In *Drosophila melanogaster,* Dicer-1 (dmDcr1) specializes in pre-miRNA processing but helicase function is not required^31^ while *Drosophila melanogaster* Dicer-2 (dmDcr2), the second Dicer in fruit flies, uses its helicase domain to recognize and bind viral and long endogenous dsRNAs^17,27,28, 32–34^ (Figure 1B). Once bound, these dsRNAs are threaded by the helicase domain to the RNaseIII sites, using the energy of ATP hydrolysis for processive cleavage into siRNA products. This processive mechanism ensures that multiple siRNAs are produced from a single dsRNA^27,32^. Another invertebrate Dicer, *Caenorhabditis elegans* Dicer-1 (ceDCR-1), likewise requires a functional helicase domain to process long endogenous/viral dsRNAs, but like dmDcr1, it also processes pre-miRNA without a requirement for ATP^13,29^. In contrast, *Homo sapiens* Dicer (hsDcr) has only been found to function in an ATP-independent manner, using its platform/PAZ domain to bind pre-miRNAs which are then distributively cleaved into mature miRNAs (Figure 1C)^24,35^. Accordingly, the role of the single mammalian Dicer enzyme in antiviral defense is controversial, as sensing of cytosolic viral dsRNAs is primarily mediated by RIG-I-Like receptors (RLRs), a family of enzymes that contain a related helicase domain^36–40^. Pathogenic dsRNA recognition by RLRs leads to production of interferon, which in turn triggers production of multiple antiviral proteins to suppress viral replication^36,41^. Thus, helicase function in invertebrate Dicers correlates with a role in antiviral defense that seems to have been replaced by RLRs in mammals.

To understand the biochemical basis of the functional diversity between Dicer helicase domains, and by inference, their roles in antiviral defense, we used phylogenetics to reconstruct evolution of Dicer’s helicase domain in animals. We included DUF283 in our analyses as recent Dicer structures reveal its role in binding dsRNA in concert with the helicase domain^17,27^. Combining phylogenetic tree construction, ancestral protein reconstruction (APR), and biochemical analyses of reconstructed proteins, we show that basal and dsRNA-dependent ATPase function was present in ancestral animal Dicer, but this capability was lost at the onset of vertebrate evolution. We also find that ancient Dicer helicases bind dsRNA more tightly than more modern and extant Dicer helicase domains. dsRNA binding to ancestral Dicer helicases stimulates ATP hydrolysis primarily by increasing the helicase domain’s affinity for ATP, as reflected in differences in K_M_ values observed in the absence and presence of dsRNA. Between the ancestor of deuterostomes and the ancestor of vertebrates, Dicer’s helicase domain lost its ability to hydrolyze ATP, while its dsRNA-binding capability declines even further. Finally, we partially resurrect ATPase function in the ancestral vertebrate Dicer helicase domain and find that loss of ATPase function is driven by amino acid substitutions distant from the catalytic pocket. We speculate that the unique role of RLRs in the interferon signaling pathway in vertebrates, and possible competition with Dicer’s helicase for the same viral dsRNAs, created an incentive to jettison an active helicase in vertebrate Dicer.

## Results

### Phylogenetic analysis of Dicer’s helicase domain reveals an ancient gene duplication of animal Dicer

Dicer’s large size (∼220 kDa) and the significant sequence divergence between its homologs and paralogs (e.g., ∼25% identity between hsDcr and dmDcr2), introduces uncertainty into multiple sequence alignments (MSAs), phylogeny construction, and ancestral protein resurrection^1,28^. Here we focused our phylogenetic analyses on the helicase domain and DUF283 (HEL-DUF), two domains involved in functional diversity across metazoan Dicers (Figure 1A). Animal Dicer sequences were retrieved from NCBI databases and truncated to leave the helicase domain and DUF283. We used the HEL-DUF sequence alignment to infer a maximum-likelihood phylogenetic tree and reconstructed the ancestral amino acid sequence on nodes from the tree. The maximum-likelihood tree revealed an early gene duplication event for HEL-DUF, where one animal Dicer (AncD1D2) was split into two major Dicer clades, AncD1 and AncD2 (Figure 1D, Figure 1-figure supplement 1). AncD1 contains the ubiquitous Dicer-1 found in most animal species, while AncD2 contains the arthropod-specific Dicer-2. The observed gene duplication is consistent with previously reported phylogenetic analyses of full-length Dicer, suggesting that the HEL-DUF region contained enough phylogenetic signal to recapitulate the broad patterns of Dicer evolution^1,2,42^.

Constraining the maximum-likelihood tree to known species relationships (Figure 1-figure supplement 2A), caused a few changes in the reconstructed sequences for AncD1D2 and AncD1_DEUT_, but the variable amino acids are not expected to significantly affect the observed biochemical properties (96% identity between HEL-DUF tree and species tree reconstruction) (Figure 1-figure supplement 3A, B). Moreover, APR using the constrained species tree did not alter the AncD1_VERT_ amino acid sequence. Furthermore, constraining arthropod Dicer-2 to a recent arthropod-specific duplication produces a more parsimonious tree that has been reported to have significantly worse likelihood scores, indicating that this tree is less likely to represent the Dicer’s true phylogeny (Figure 1-figure supplement 2B)^1^.

Select nodes in the maximum likelihood tree were subjected to APR using RAXML-NG. Amino acid sequences for AncD1D2, AncD2_ARTH_, AncD1_ARTH/LOPH/DEUT_, AncD1_LOPH/DEUT_, AncD1_DEUT_ and AncD1_VERT_, were predicted with a high degree of confidence. AncD1_VERT_ had more than 95% of sites with posterior probabilities of 0.8 or above, while older constructs had an average of 75% of sites with posterior probabilities above 0.8 (Figure 1-figure supplement 4A). Ancestral constructs were expressed recombinantly using baculovirus in Sf9 insect cells and purified to 99% homogeneity (Figure 1-figure supplement 4B)^43^. Protein identity was confirmed with LC/MS/MS.

### Ancient animal Dicer helicase domain possessed dsRNA-stimulated ATPase function

Certain extant Dicer enzymes have ATP hydrolysis activity, while others do not, suggesting either a gain or loss of this activity during evolution. To understand the source of this variation, we performed multiple-turnover ATP hydrolysis assays of ancestral HEL-DUF proteins with and without dsRNA. We observed ATP hydrolysis in the most recent common ancestor of hsDcr and dmDcr2, AncD1D2 (Figure 2A), leading to the important conclusion that extant animal Dicers with no dependence on ATP, such as hsDcr, lost the capacity for ATP hydrolysis subsequent to this period in animal evolution. Basal ATP hydrolysis activity was present at low levels in AncD1D2 (k_obs_: 0.06 min^-1^) and was improved upon addition of dsRNA (Figure 2A, Table 1).

**Figure 2.**
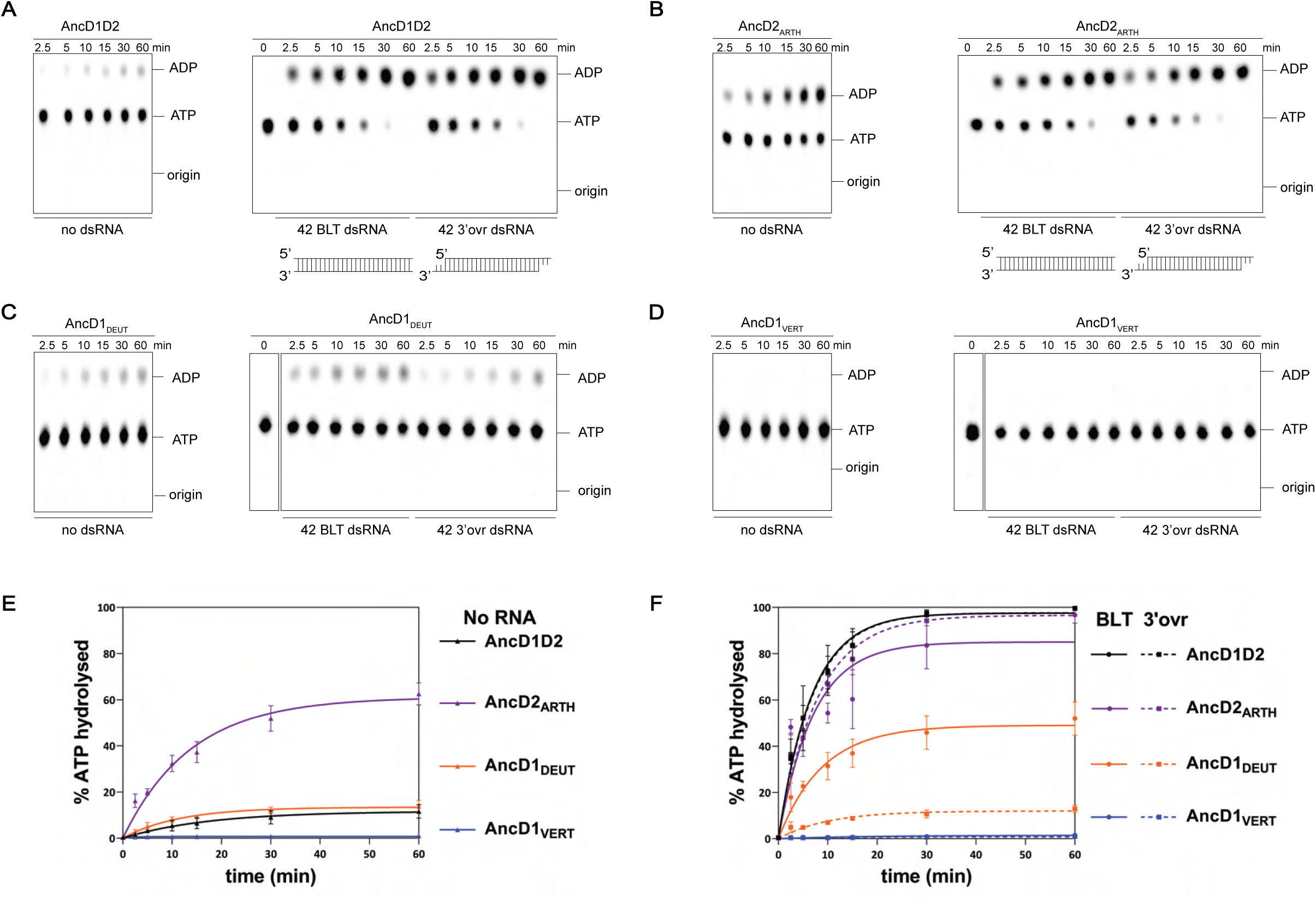
ATP hydrolysis capability is present in ancestral metazoan Dicer but lost at the common ancestor of vertebrates. **A-D.** PhosphorImages of representative Thin-layer Chromatography (TLC) plates showing hydrolysis of 100µM ATP (spiked with α-^32^P-ATP) by 200nM ancestral HEL-DUFs for various times as indicated, at 37°C, in absence of dsRNA or in the presence of 400nM 42 base-pair dsRNA with BLT or 3’ovr termini (see cartoons; not radiolabeled). **E.** Graph shows quantification of ATP hydrolysis assays (A-D) performed with select ancestral HEL-DUF enzymes in the absence of dsRNA. Data for “NO RNA” reactions were fit to the pseudo-first order equation y = y_o_ + A x (1-e^-kt^); where y = product formed (ADP in µM); A = amplitude of the rate curve, y_o_ = baseline (∼0), k = pseudo-first-order rate constant = k_obs_; t = time. Data points are mean ± SD (n≥3). **F.** Graph shows quantification of ATP hydrolysis assays (A-D) performed with select ancestral HEL-DUF enzymes in the presence of dsRNA. Reactions with RNA were fit in two phases, first a linear phase for data below the first timepoint at 2.5 minutes, then a pseudo-first order exponential equation for remaining data. Equation, y = y_o_ + A x (1-e^-kt^); where y = product formed (ADP in µM); A = amplitude of the rate curve, y_o_ = baseline (∼0), k = pseudo-first-order rate constant = k_obs_; t = time. Data points are mean ± SD (n≥3).

**Table 1:** Summary of kinetic data for ATP hydrolysis with 100µM ATP.

| Construct | $k_{burst}$ (μM/min), NO RNA<br>$k_{obs}$ (min<sup>-1</sup>) | $k_{burst}$ (μM/min), BLT dsRNA<br>$k_{obs}$ (min<sup>-1</sup>) | $k_{burst}$ (μM/min), 3'ovr dsRNA<br>$k_{obs}$ (min<sup>-1</sup>) |
| --- | --- | --- | --- |
| AncD1D2 | -<br>0.06 ± 0.01 | 14.3 ± 1.7<br>0.11 ± 0.03 | 13.9 ± 0.5<br>0.11 ± 0.02 |
| AncD2<sub>ARTH</sub> | 6.47 ± 0.8<br>0.05 ± 0.01 | 19.3 ± 0.9<br>0.04 ± 0.02 | 14.6 ± 2.3<br>0.08 ± 0.02 |
| AncD1<sub>ARTH/LOPH/DEUT</sub> | -<br>0.01 ± 0.01 | 25.1 ± 0.7<br>0.41 ± 0.02 | 16.0 ± 3.8<br>0.21 ± 0.04 |
| AncD1<sub>LOPH/DEUT</sub> | -<br>0.07 ± 0.02 | 7.4 ± 0.3<br>0.04 ± 0.01 | 3.8 ± 0.6<br>0.03 ± 0.01 |
| AncD1<sub>DEUT</sub> | -<br>0.09 ± 0.02 | 6.0 ± 0.7<br>0.06 ± 0.03 | 1.4 ± 0.03<br>0.06 ± 0.02 |
| AncD1<sub>VERT</sub> | - | - | - |

Adding dsRNA to AncD1D2 showed a dramatic increase in the ADP produced over time (Figure 2A, right panel). To enable better comparison of efficient HEL-DUF ancestors, and minimize effects of substrate depletion during the reaction, data were modeled as a two-phase exponential curve. The first phase was represented by a fast linear burst of ATP hydrolysis capturing a transient zero order reaction where rate is independent of ATP concentration, and the second phase was modeled as a slow, first order exponential increase in ATP hydrolysis, where, because of robust hydrolysis, ATP concentration falls below some affinity threshold for the enzyme (Figure 2E, F, Table 1)^44^. Without dsRNA, ATP hydrolysis is a slow first order reaction because the concentration of ATP in this reaction (100µM) is likely orders of magnitude below the affinity threshold. Addition of dsRNA with BLT termini to AncD1D2, a substrate designed to mimic termini of certain RNA viruses, promoted hydrolysis of ATP in the fast phase (k_burst:_ 14.3 µM/min) as well as doubling the rate constant of the slow phase (k_obs_: 0.11 min^-1^) (Table 1). Similar rates were observed when a dsRNA with a 2-nucleotide (nt) 3’ overhang (3’ovr) (pre-miRNA mimic), was used (Table 1). Lack of dsRNA terminus discrimination suggests a substrate promiscuity that is absent in modern Dicers, where BLT dsRNA is the optimal substrate for the helicase domain^28–30,32^.

The arthropod Dicer-2 ancestor, AncD2_ARTH_, was more efficient at hydrolyzing ATP in the absence of dsRNA than AncD1D2 and all other ancestors tested (Figure 2B, E), showing a two-phase reaction resembling a dsRNA-stimulated reaction (k_burst_: 6.47 µM/min, k_obs_: 0.05 min-1; Table 1). This efficient hydrolysis in the absence of dsRNA suggests that AncD2_ARTH_ refined its ATP binding pocket to produce high affinity for ATP even in the absence of nucleic acid. Addition of BLT dsRNA increased the rate of the fast phase (k_burst_: 19.3 µM/min), with slightly better efficiency compared to 3’ovr dsRNA (k_burst_:14.6 µM/min) (Table 1, Figure 2B, F), perhaps foreshadowing the terminus discrimination seen in modern dmDcr2^30,32^. Interestingly, terminus discrimination was also observed in Dicer-1 ancestors, suggesting the foundation of this discrimination existed in AncD1D2, even if not observable at the conditions tested. The deuterostome Dicer-1 node immediately preceding vertebrate Dicer-1, AncD1_DEUT_, had a pattern of basal and dsRNA-stimulated ATP hydrolysis similar to AncD1D2 (Table 1) (Figure 2C, E, F) but had obvious differences in the efficiency of hydrolysis triggered by different dsRNA termini (BLT, k_burst_: 6.0 µM/min; 3’ovr, k_burst_: 1.4 µM/min) (Table 1).

Most importantly, in the ancestor of vertebrate Dicer, AncD1_VERT_, ATP hydrolysis was undetectable (Figure 2D), consistent with observations that modern hsDcr is incapable of ATP hydrolysis^35^, and indicating that loss of Dicer ATPase function is a vertebrate-wide phenomenon, driven by evolutionary events between the deuterostome and vertebrate ancestor. This period in evolution coincides with whole genome duplications important for vertebrate evolution, the expansion of the miRNA repertoire and their role in gene regulation, and most critically, the advent of the interferon molecule and its role in innate immunity^45–48^.

### Ancestral HEL-DUF binds dsRNA with higher affinity than modern Dicer HEL-DUF

Our experiments indicated that dsRNA improves ATP hydrolysis by ancient Dicer helicase domains, in some cases in a terminus-dependent manner. We wondered if terminus discrimination occurred during initial dsRNA binding. To investigate the dsRNA•HEL-DUF interaction, as well as how it is affected by ATP, we used electrophoretic mobility shift assays (EMSAs) with HEL-DUFs to measure the dissociation constant (K_d_) in the presence and absence of ATP, using BLT or 3’ovr dsRNA (Figure 3A)^49,50^.

**Figure 3.**
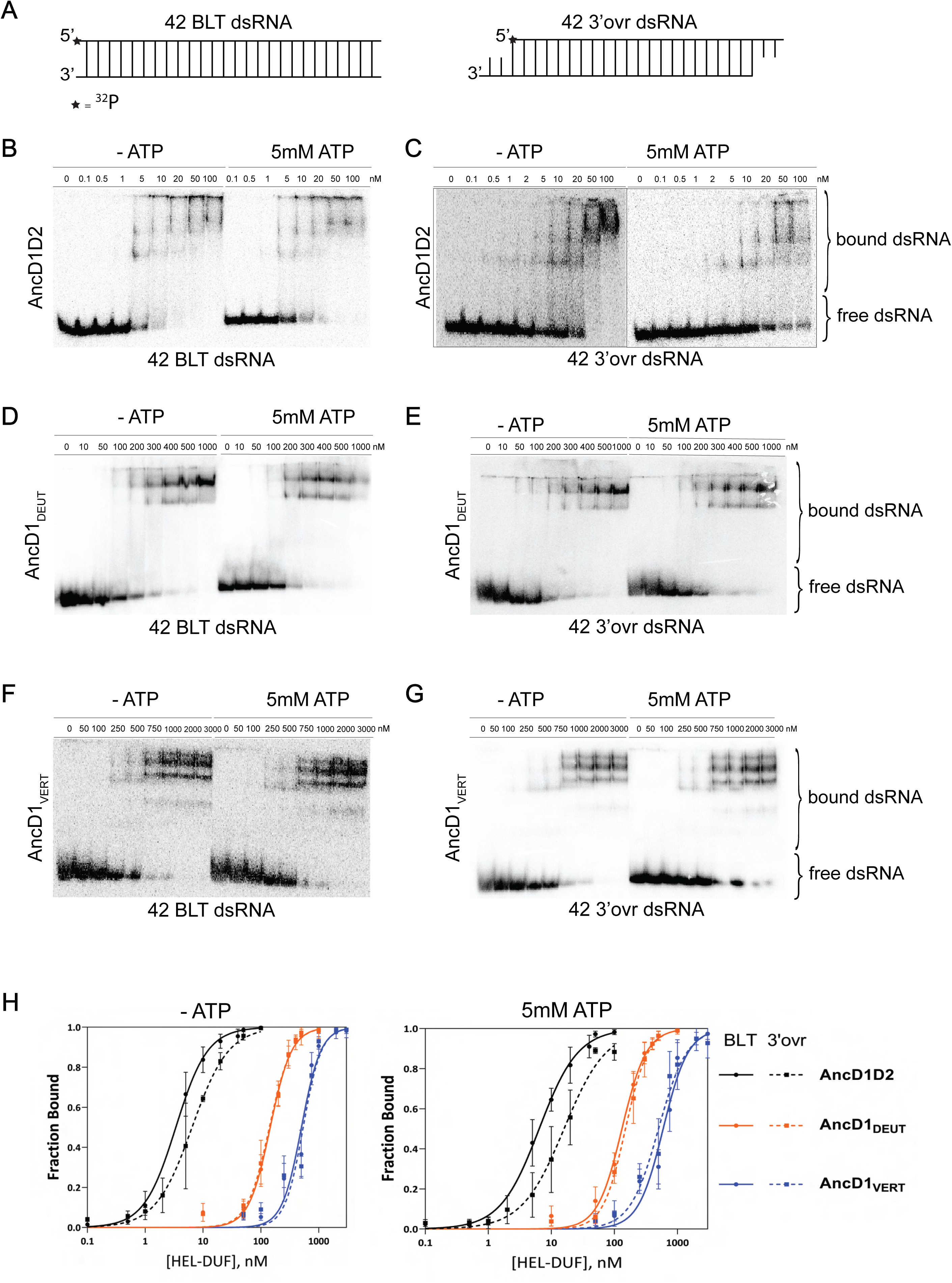
Binding affinity of ancestral HEL-DUF proteins to BLT and 3’ovr dsRNA in the presence and absence of ATP. **A.** Cartoon of dsRNAs used in (B-G) showing position of 5’ ^32^P (*) on top, sense strand. **B-G.** Representative PhosphorImages showing gel mobility shift assays using select ancestral HEL-DUF constructs, as indicated, and 42-basepair BLT or 3’ovr dsRNA in the absence (-) or presence of 5mM ATP at 4°C. **H.** Radioactivity in PhosphorImages as in A-G was quantified to generate binding isotherms for ancestral HEL-DUF proteins. Fraction bound was determined using radioactivity for dsRNA_free_ and dsRNA_bound_. Data were fit to calculate dissociation constant, K_d_, using the Hill formalism, where fraction bound = 1/(1 + (K_d_ /[P])). Data points, mean ± SD (n≥3).

All ancestral proteins bound dsRNA and showed multiple shifts that typically decreased in mobility with increasing protein concentration. AncD1D2, the most ancient construct tested, displayed tight binding to BLT dsRNA without ATP (K_d_, 3.4nM), while the addition of 5mM ATP caused a 2-fold reduction in affinity (K_d_, 6.4nM; Table 2, Figure 3B, H). Binding to 3’ovr dsRNA was similarly tight, albeit with an ∼2-fold reduction in binding affinity compared to BLT in the absence or presence of ATP (Table 2, Figure 3C, H). This suggests that dsRNA binding is the earliest point of terminus discrimination for ATP hydrolysis with ancestral HEL-DUFs. Possibly, the observed lower affinity with ATP occurs because ATP hydrolysis promotes dissociation, or translocation that results in HEL-DUF sliding off dsRNA.

**Table 2:** Dissociation constants for dsRNA binding to ancestral HEL-DUFs.

| Construct | $K_d$ (nM) BLT, NO ATP<br>Hill coefficient | $K_d$ (nM) 3'ovr, NO ATP<br>Hill coefficient | $K_d$ (nM) BLT, 5mM ATP<br>Hill coefficient | $K_d$ (nM) 3'ovr, 5mM ATP<br>Hill coefficient |
| --- | --- | --- | --- | --- |
| AncD1D2 | 3.4 ± 0.4<br>1.6 ± 0.2 | 6.5 ± 0.8<br>1.4 ± 0.2 | 6.4 ± 0.7<br>1.4 ± 0.2 | 15.9 ± 2.4<br>1.3 ± 0.2 |
| AncD2<sub>ARTH</sub> | n.d. | n.d. | n.d. | n.d. |
| AncD1<sub>ARTH/LOPH/DEUT</sub> | 23.8 ± 2.2<br>2.0 ± 0.4 | 40.1 ± 3.7<br>1.7 ± 0.3 | 17.5 ± 2.1<br>1.6 ± 0.3 | 17.3 ± 1.5<br>1.7 ± 0.2 |
| AncD1<sub>LOPH/DEUT</sub> | 60.9 ± 5.9<br>1.9 ± 0.3 | 90.8 ± 8.8<br>1.4 ± 0.2 | 38.0 ± 3.5<br>2.3 ± 0.5 | 49.0 ± 5.2<br>1.4 ± 0.2 |
| AncD1<sub>DEUT</sub> | 145.1 ± 9.1<br>2.3 ± 0.3 | 140.0 ± 8.9<br>2.3 ± 0.3 | 131.8 ± 8.7<br>2.2 ± 0.3 | 149.8 ± 8.3<br>2.4 ± 0.3 |
| AncD1<sub>VERT</sub> | 502.4 ± 40.5<br>2.6 ± 0.5 | 537.8 ± 47.5<br>2.8 ± 0.6 | 592 ± 48.6<br>2.2 ± 0.4 | 500.3 ± 58.0<br>1.9 ± 0.4 |

An obvious feature evident from the binding isotherms (Figure 3H) was that the more ancient the HEL-DUF construct in our tree, the tighter its interaction with dsRNA, regardless of the presence or absence of ATP. Comparison of K_d_ values (Table 2) revealed other interesting trends. AncD1_DEUT_, the common ancestor of deuterostomes, which include humans, had a lower binding affinity for dsRNA compared to AncD1D2, regardless of termini or the presence of ATP (∼10-50-fold, Table 2, Figure 3D, H). In addition, the ability to distinguish BLT and 3’ovr dsRNA largely disappeared, and ATP had little effect on dsRNA binding (Table 2, Figure 3E, H). This absence of discrimination between termini or ATP-bound states stands in contrast to observations for AncD1D2 and extant *Drosophila melanogaster*, whose helicase domains bind BLT dsRNA better than 3’ovr dsRNA^28,33^. Also, this lack of discrimination in binding does not match the sensitivity to dsRNA termini observed in ATP hydrolysis (Figure 2C), suggesting an additional discriminatory step exists between initial dsRNA binding and ATP hydrolysis. Another possibility is that for some constructs but not for others, ATP’s effect on dsRNA binding is muted or altered at the low temperatures (4°C) where EMSAs were performed.

Binding of dsRNA to AncD1_VERT_ resembled binding to AncD1_DEUT_ in that affinity did not depend on termini or ATP (Table 2, Figure 3F, G, H). However, AncD1_VERT_ binding to dsRNA was weaker across all conditions with ∼4.5-fold reduction in affinity compared to AncD1_DEUT_, and ∼30-150-fold reduction compared to AncD1D2 (Table 2). Interestingly, the K_d_ values measured for AncD1_VERT_ were similar to values reported for the modern hsDcr helicase domain, or hsDcr with the platform/PAZ domain mutated to abolish competing binding, hinting that this construct correlates with extant biology^51,52^. One possibility is that during evolution, vertebrate HEL-DUF’s affinity decreased as the platform/PAZ domain began to play a more significant role in binding 3’ovr termini of pre-miRNAs, and RLRs co-opted binding of virus-like BLT dsRNAs. In summary, the more ancient the HEL-DUF construct in our tree, the tighter the dsRNA•HEL-DUF interaction, with the deuterostome HEL-DUF ancestor losing the ability to discriminate dsRNA termini by binding.

### BLT dsRNA improves ATP hydrolysis by improving the association of ATP to HEL-DUF

Our analyses so far showed that dsRNA markedly altered the kinetics and efficiency of ATP hydrolysis. To understand how dsRNA binding affected the interaction of the helicase with ATP, we performed Michaelis-Menten analyses. We focused on determining kinetics for ATP hydrolysis catalyzed by AncD1D2 and AncD1_DEUT_, to gain information about two Dicer-1 ancestors separated by vast evolutionary distance. Without added dsRNA, basal ATP hydrolysis for AncD1D2 had a k_cat_ of 1117 min^-1^ and a K_M_ of 35.8mM (Table 3, Figure 4A, Figure 4-figure supplement 1). Adding excess BLT dsRNA caused k_cat_ to drop ∼8 fold to 147.8 min^-1^ while K_M_ dropped ∼140-fold to 0.26mM, indicating that although binding of BLT dsRNA to the AncD1D2 causes a reduction in the ATP turnover efficiency, it concurrently triggers a tighter association with ATP, leading to ∼19-fold net improvement in k_cat_/K_M_ (Table 3). This improvement in efficiency is primarily evident at lower ATP concentrations that fall in the range of intracellular ATP concentrations^53^. (Figure 4A, right panel). These observations also explain the appearance of the fast linear phase in the multiple-turnover hydrolysis reactions performed with 100µM ATP (Table 1, Figure 2). At this “lower” ATP concentration, we speculate that dsRNA binding causes a conformational change in the helicase domain that allows tighter association with ATP, enabling a more efficient hydrolysis reaction, until ATP concentration falls below a threshold where the reaction slows. Without dsRNA, only slow ATP hydrolysis is available as the helicase rarely samples the conformations that allow tight interactions with ATP.

**Figure 4.**
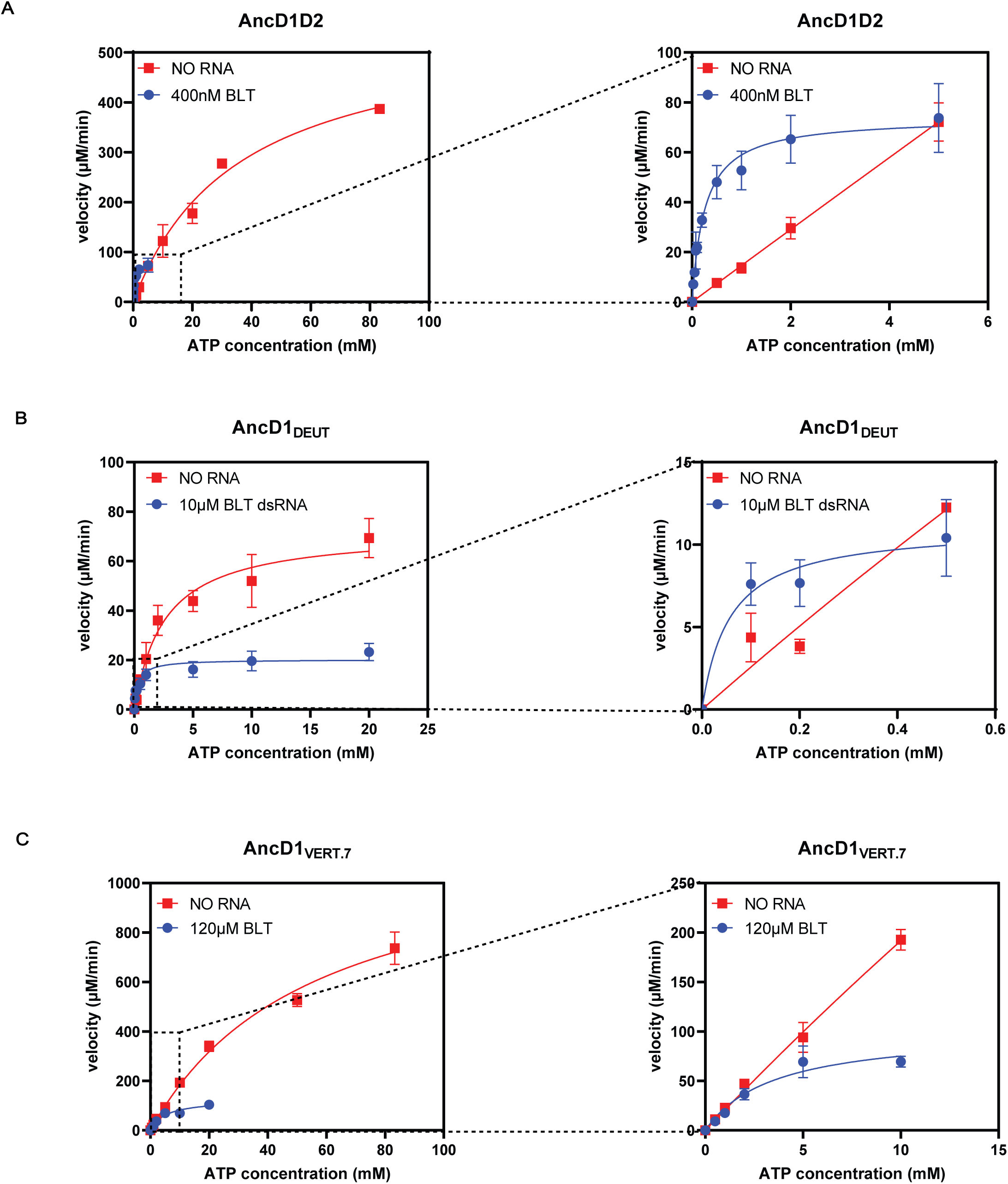
BLT dsRNA improves efficiency of ATP hydrolysis by improving affinity of ATP to ancient HEL-DUF enzymes. **A.** Michaelis-Menten plots for basal and dsRNA-stimulated ATP hydrolysis by AncD1D2. Basal ATP hydrolysis measured at 500nM AncD1D2, while dsRNA-stimulated hydrolysis is measured at 100nM. Velocities for dsRNA-stimulated reaction have been multiplied by 5 to normalize this concentration difference. Right: inset showing Michaelis-Menten plot at low ATP concentrations. Hydrolysis data for individual ATP concentrations is included in Figure 4-figure supplement 1. **B.** Michaelis-Menten plots for basal and dsRNA-stimulated ATP hydrolysis by 500nM AncD1_DEUT_. Right: inset showing Michaelis-Menten plot at low ATP concentrations. Hydrolysis data for individual ATP concentrations is included in Figure 4-figure supplement 1. **C.** Michaelis-Menten plots for basal and dsRNA-stimulated ATP hydrolysis by 5µM AncD1_VERT.7_. Right: inset showing Michaelis-Menten plot at low ATP concentrations. Hydrolysis data for individual ATP concentrations is included in Figure 4-figure supplement 1. Data points, mean ± SD (n≥3).

**Table 3:** Michaelis-Menten parameters for steady state ATP hydrolysis reactions.

|  |  |  |  |
| --- | --- | --- | --- |
| Construct | $k_{cat}$ ($\text{min}^{-1}$) | $K_M$ ($\mu\text{M}$) | $k_{cat}/K_M$ ($\mu\text{M}^{-1} \text{min}^{-1}$) |
| AncD1D2, no dsRNA | $1117 \pm 94.5$ | $35812 \pm 6367$ | 0.031 |
| AncD1D2, BLT dsRNA | $147.8 \pm 6.3$ | $256 \pm 67.5$ | 0.577 |
| AncD1<sub>DEUT</sub>, no dsRNA | $144.1 \pm 14.9$ | $2550 \pm 855$ | 0.055 |
| AncD1<sub>DEUT</sub>, BLT dsRNA | $40.31 \pm 3.78$ | $336.4 \pm 142$ | 0.12 |
| AncD1<sub>VERT</sub>, no dsRNA | - | - | - |
| AncD1<sub>VERT.7</sub>, no dsRNA | $257.7 \pm 28.7$ | $61739 \pm 12886$ | 0.004 |
| AncD1<sub>VERT.7</sub>, BLT dsRNA | $24.87 \pm 3.52$ | $5173 \pm 1929$ | 0.005 |

For the AncD1_DEUT_ construct, basal hydrolysis proceeded with a k_cat_ of 144.1 min^-1^, ∼8-fold lower than the rate recorded for the AncD1D2 reaction (Table 3, Figure 4B, Figure 4-figure supplement 1C). Association of ATP with the AncD1_DEUT_ construct, as measured by K_M_, was ∼14-fold tighter leading to similar k_cat_/K_M_ values for both enzymes in the absence of dsRNA (Table 3). Adding BLT dsRNA caused a reduction in k_cat_ by a factor of ∼4, while reducing K_M_ by a factor of ∼8 to give a 2-fold net increase in k_cat_/K_M_ (Table 3, Figure 4B, Figure 4-figure supplement 1D). As with the AncD1D2 construct, improvement is primarily mediated by improved association of enzyme with ATP, evident at lower ATP concentrations (Figure 4B, right panel). Thus, we observed a trend where saturation of the helicase domain with BLT dsRNA caused improved ATP association to the catalytic ATPase site.

### Reversing historical substitutions in AncD1_VERT_ partially rescued ATP hydrolysis

To acquire insight into the reason for loss of function in the ancestor of vertebrate Dicer, AncD1_VERT_, we compared Michaelis-Menten kinetics of AncD1_VERT_ and AncD1_DEUT_. However, even at higher protein concentrations (5µM), it was impossible to confidently discern whether AncD1_VERT_ produced a signal over background using our TLC-based assay (data not shown). Thus, we compared amino acid sequences of ancestral HEL-DUFs that retained ATPase activity to the sequence of the inactive vertebrate HEL-DUF ancestor, and identified substitutions that might be responsible for the loss of activity (Figure 4-figure supplement 2). ATP hydrolysis and dsRNA-binding data for the gene tree-species tree incongruent nodes, AncD1_ARTH/LOPH/DEUT_ and AncD1_LOPH/DEUT_, proved useful here, allowing a deeper analysis of sequence-function relationships of Dicer’s helicase domain (Figure 4-figure supplement 3, 4). We created variants of AncD1_VERT_, each with a subset of these substitutions, and purified two constructs, AncD1_VERT.1_ and AncD1_VERT.7_ (Figure 4-figure supplement 2). In AncD1_VERT.1_, we reverted 20 amino acids which were either close to ATP binding/hydrolysis amino acid residues or to the ATPase active site in the tertiary structure, but this construct remained devoid of ATPase activity (data not shown). However, in AncD1_VERT.7_, a construct containing an additional 21 amino acid substitutions distant from the catalytic site, ATPase activity was rescued, and we measured its Michaelis-Menten kinetics. Basal ATP hydrolysis had a k_cat_ of 257.7 min^-1^ and a K_M_ of 61.7mM (Table 3, Figure 4C, Figure 4-figure supplement 1E). Enzyme turnover was more efficient than AncD1_DEUT_, but less efficient than AncD1D2, suggesting a rescue of the enzyme’s inherent catalytic activity. However, the high K_M_ value indicated that AncD1_VERT.7_ was not rescued for a tight association with ATP, and the k_cat_/K_M_ value for this construct was ∼10-fold lower than k_cat_/K_M_ for both AncD1_DEUT_ and AncD1D2 (Table 3). Adding BLT dsRNA caused k_cat_ to drop ∼10-fold to 24.9min^-1^ while improving the K_M_ by ∼12-fold, therefore yielding no net improvement in k_cat_/K_M_. (Table 3, Figure 4C, Figure 4-figure supplement 1F). Our observations indicate that we have partially rescued ATPase activity in vertebrate HEL-DUF as well as the conformational changes that occur upon dsRNA binding. AncD1_VERT.1_, constructed by reversing candidate amino acid substitutions close in proximity to the conserved ATPase motifs, did not rescue ATPase activity. Instead, we found that amino acids distant from the ATP binding pocket were essential for resurrecting ATP hydrolysis in the vertebrate ancestor of Dicer.

## Discussion

Phylogenetic tools have been used to analyze the evolution of the platform/PAZ domain in plant and animal Dicers (Figure 1A), shedding light on one source of functional diversity in eukaryote Dicer^1^. Here we focused on evolution of Dicer’s helicase domain in animals. An ATP-dependent helicase domain is important for Dicer’s antiviral role in invertebrates such as dmDcr2 and ceDCR-1, while mammalian Dicer has not been observed to require ATP^29,54–56^. One model is that mammalian Dicer’s helicase domain exists to stabilize the interaction of pre-miRNAs with the platform/PAZ domain during processing to mature miRNAs^24,26,57^. Arthropods and nematodes are invertebrate ecdysozoan protostomes, and so far, these are the only two phyla where Dicer’s helicase domain is known to be essential for antiviral defense. Is this property specific to Ecdysozoa or is it more widespread among other invertebrates and bilaterians? Why is it absent in mammalian Dicer? The catalytic motif in this family of helicases is the DECH box, also known as motif II^58,59^. The DECH box is conserved between arthropods, nematodes and mammals but significant divergence in amino acid sequence of hsDcr, dmDcr2, and ceDCR-1, makes it challenging to answer these questions simply by analyzing amino acid variation. By performing APR, we generated evolutionary intermediates that revealed more subtle changes in amino acid variation and biochemical function, allowing insight into the biochemical properties of the ancient helicase domain and how these evolved to give rise to extant Dicer’s roles in gene regulation and antiviral defense. The robust sequence-function analysis of Dicer’s helicase domain provided by APR also allowed us to probe a larger sequence space that may not exist in any ancient or extant Dicers, as is the case for gene tree-species tree incongruent nodes like AncD1_ARTH/LOPH/DEUT_ and AncD1_LOPH/DEUT_.

### Ancestral animal Dicer possessed an active helicase domain

Our analysis revealed that AncD1D2, the common ancestor of dmDcr2, hsDcr, and ceDCR-1, retained the capability to hydrolyze ATP (Figure 2A). In addition, our phylogeny construction, performed with the helicase domain and DUF283, recapitulates the early gene duplication event reported previously in phylogenetic studies of full-length Dicers (Figure 1D)^1,2^. Plants and fungi have been reported to have Dicer or Dicer-like proteins with active helicase domains^4,17^ so it stands to reason that early animal Dicer descended from an ancestral eukaryotic Dicer with an active helicase^60^. The ATP hydrolysis observed in our Dicer ancestors are predicted to be coupled to some motor function as observed in extant arthropod Dicer-2. Future studies will determine if these constructs couple ATP hydrolysis to translocation and/or unwinding like dmDcr2, or to some other function like terminus discrimination.

### Dicer ATPase function is lost at the onset of vertebrate evolution

As animals evolved from deuterostomes (AncD1_DEUT_) to vertebrates (AncD1_VERT_), Dicer lost the ability to hydrolyze ATP (Figure 2C, D). The loss of both intrinsic and dsRNA-stimulated ATPase activity between AncD1_DEUT_ and AncD1_VERT_ can be attributed to one of a number of evolutionary events. Whole genome duplication events that occurred at the onset of vertebrate evolution may have caused subfunctionalization of Dicer’s helicase domain, as other antiviral sensors like RIG-I-like receptors (RLRs) and Toll-like-receptors (TLRS) co-opted the role of sensing pathogen-associated molecular patterns (PAMPs)^45,61^. Upon binding dsRNAs, these receptors trigger an enzyme cascade that ultimately produces interferon, a molecule that came into being at the onset of vertebrate evolution^46^. In support of this model, there are multiple examples of antagonism between the RNA interference (RNAi) pathway and the RLR signaling pathway in mammals^40,62–64^. The model is further supported by our dsRNA binding studies where the observed weak dsRNA binding by the vertebrate helicase domain suggests that cytosolic dsRNA recognition was taken over by RLRs (Figure 3, Table 2). If this model is true, it raises new questions. Is vertebrate Dicer’s loss of ATPase function a consequence of the RLR-interferon axis, a protein-based antiviral system, taking over antiviral defense from RNAi, a nucleic-acid based defense mechanism? Could vertebrates maintain both modes of antiviral defense if loss of function in Dicer’s helicase domain was reversed? Further studies on the selection pressures exerted on Dicer in different species are required to answer these questions.

### Changes in Dicer helicase domain’s conformation influence ATPase activity

The effects of dsRNA binding on ancestral Dicer helicase domains show parallels to previously reported results for extant Dicers. In the absence of dsRNA, hsDcr and dmDcr2 helicase domains primarily exist in an “open” conformation, with DUF283 wedged behind the Hel1 subdomain (Figure 5, A and B)^24,27^. dsRNA binding to dmDcr2’s helicase domain causes a conformational change that brings DUF283 to the cleft of the helicase domain to cap the dsRNA substrate (Figure 5C), while Hel2 and Hel2i shift their relative positions to create a “closed” conformation. In the closed conformation, the distance between the “DECH” box in Hel1 (Motif II) and the arginine finger motif (Motif VI) in Hel2 is reduced from 13. 28Å to 4.26 Å (Figure 5D)^27^. In helicases formed by two RecA domains, the shorter distance is predicted to be bridged by water, the attacking nucleophile for cleavage of the gamma-phosphate of the ATP molecule^65,66^. This change in conformation is consistent with the reduction in the K_M_ values for ATP when BLT dsRNA is included in the ATP hydrolysis reaction for AncD1D2 and AncD1_DEUT_ (Figure 4, Table 3). Along the same lines, a K_M_ of 14µM was reported for the dmDcr2 ATPase reaction in the presence of BLT dsRNA^15^. We predict that excluding dsRNA would also reduce dmDcr2’s affinity for ATP, explaining why it does not hydrolyze ATP in the absence of dsRNA^30^.

**Figure 5.**
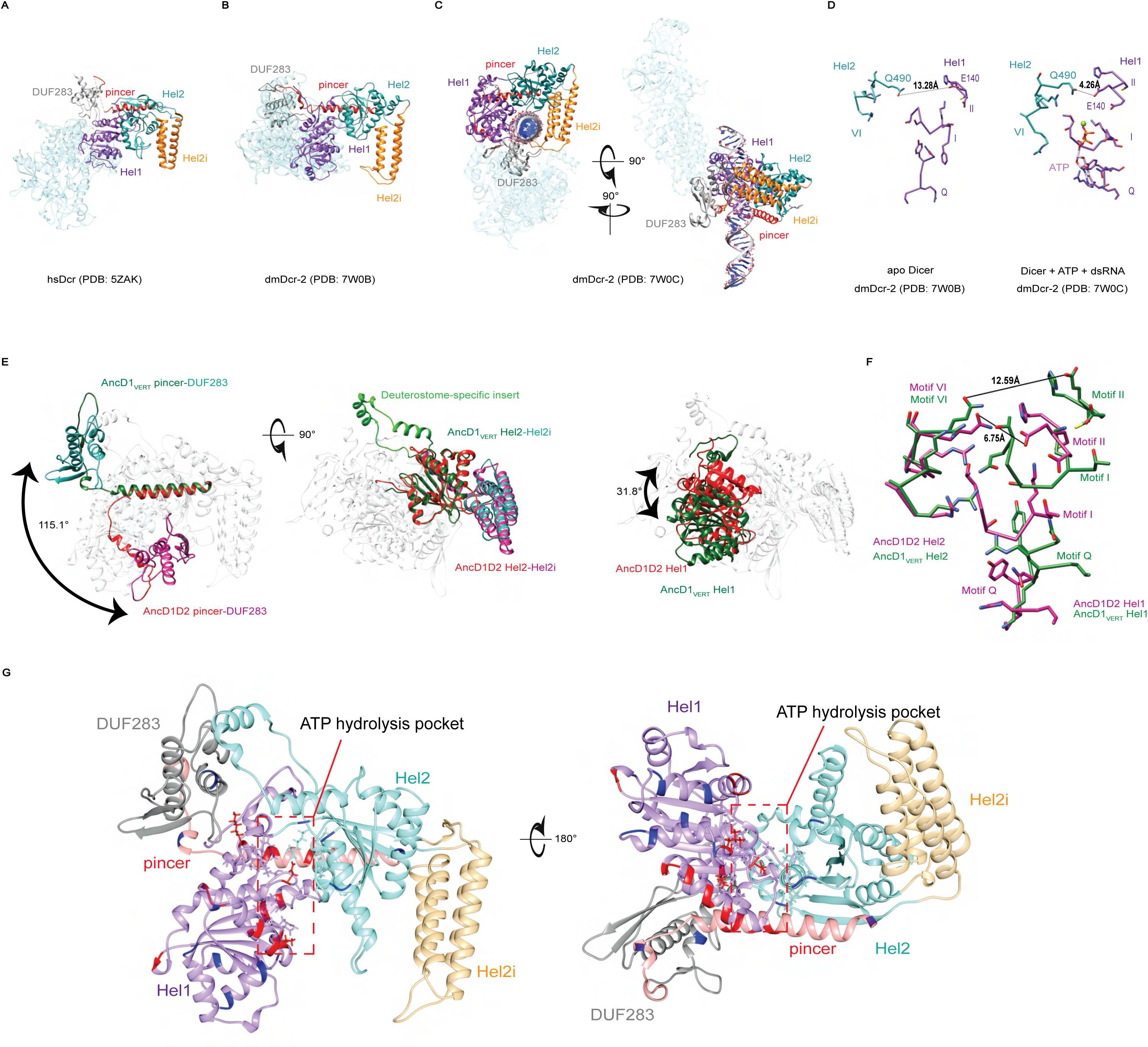
dsRNA binding triggers conformational changes in the HEL-DUF domains of Dicer. **A.** Bottom-up view of the structure of hsDcr Dicer in the apo state (PDB: 5ZAK). Helicase subdomains and DUF283 are colored. Rest of enzyme is transparent. **B.** Bottom-up view of the structure of dmDcr2 in the apo state (PDB: 7W0B). Helicase subdomains and DUF283 are colored for visibility. Rest of enzyme is transparent. **C.** Structure of dmDcr2 bound to dsRNA in the “early translocation” state (PDB: 7W0C). Helicase subdomains and DUF283 are colored for visibility. **D.** Details of interactions at the ATP binding pocket of dmDcr2, comparing the distance between Motif II and Motif VI for the apo enzyme and the enzyme in the presence of ATP and dsRNA. Green sphere is magnesium ion, a cofactor in Sf2 helicase ATP hydrolysis. **E.** Structural alignment of predicted structures for AncD1D2 and AncD1_VERT_ HEL-DUFs showing conformational differences in position of Hel1 and pincer subdomains and DUF283. Pincer and DUF293, left panel; Hel2 and Hel2i, middle panel; Hel1, right panel. Green and teal coloring represent AncD1_VERT_ subdomains, red and violet coloring represent AncD1D2 subdomains. Deuterostome-specific insert refers to a Hel2 insertion present in AncD1_DEUT_ and AncD1_VERT_. Structural predictions were performed with RosettaFold and AlphaFold2. pLDDT score: 81.76 for AncD1D2, 74.60 for AncD1_VERT_. **F.** Details of the interactions of the ATP binding pocket for AncD1D2 and AncD1_VERT_, showing a wider cleft between Motif II and Motif VI for AncD1_VERT_ (violet) compared to AncD1D2 (green). **G.** RosettaFold predicted structures for AncD1_VERT_ (transparent) showing sites of amino acid substitutions for both AncD1_VERT.1_ and AncD1_VERT.7_ marked in red, and amino acid substitutions unique to AncD1_VERT.7_ marked in blue. ATP hydrolysis pocket is depicted.

AncD1_VERT_ structures predicted by AlphaFold2 and RosettaFold have an open conformation, while AncD1D2 resembles the closed conformation (Figure 5E). All the other ATPase-competent ancestral HEL-DUFs also have a closed conformation (not shown). While these predictions are snapshots of a singular conformation from an ensemble of possible conformations, it is intriguing that the conformational differences between ancestral HEL-DUFs match our experimentally determined biochemical properties. In AncD1_VERT_, DUF283 (teal) is behind the helicase domain while AncD1D2’s DUF283 (violet) caps the helicase cleft as in the closed conformation of dmDcr2 (Figure 5E, left panel). Hel2 (green) and Hel2i (teal) subdomains of AncD1_VERT_ align closely with the corresponding subdomains in AncD1D2 (red and violet respectively) (Figure 5E, middle panel). However, AncD1_VERT_’s Hel1 (green) leans away from the Hel2-Hel2i rigid body by 31.8° compared to AncD1D2’s Hel1 subdomain (red) (Figure 5E, right panel). This difference in conformation affects the ATP binding pocket which exists at the interface between Hel1 and Hel2, showing a wider distance between helicase motif II and motif VI in AncD1_VERT_ compared to AncD1D2 (Figure 5F). This may explain why AncD1_VERT_ is incapable of basal hydrolysis even when catalytic motifs are repaired in AncD1_VERT.1_. Non-catalytic motifs that affect conformation and helicase subdomain movement mediate the loss of ATPase function in vertebrate Dicer’s helicase domain.

### ATP and dsRNA binding are limiting factors in vertebrate Dicer helicase function

The role of hsDcr in antiviral defense is controversial^10,11, 67–70^. The current consensus is that hsDcr is more relevant for antiviral defense in stem cells, while RLRs and interferon signaling predominate in somatic cells^40,64^. HsDcr’s helicase domain is, however, not involved in stem-cell specific antiviral function, and in fact, cleavage of viral or endogenous long dsRNA is improved when the helicase domain is truncated or removed^71,72^. This suggests that hsDcr’s helicase domain is incapable of coupling dsRNA translocation to ATP hydrolysis. Our dsRNA binding data reinforce this observation: dsRNA binding is significantly worse in AncD1_VERT_ than it is for AncD1D2, suggesting that vertebrates in general do not use Dicer’s helicase domain for antiviral defense (Figure 3). Reinforcing this model, cryo-EM structures of hsDcr report an open conformation for hsDcr’s helicase domain even in the presence of dsRNA (Figure 1C), and cleavage of long dsRNAs by hsDcr is mediated by direct binding to the platform/PAZ domain with no requirement for ATP^24,25^.

Canonical SF2 helicase ATP binding/hydrolysis motifs, like the eponymous “DECH” box, are conserved between hsDcr and dmDcr2, but outside these motifs, the primary sequence varies significantly^28^. Using sequence- and structure-based alignments of our ancestral HEL-DUF constructs, we identified candidate historical substitutions outside the catalytic motifs (Figure 4-figure supplement 2) that caused the loss of intrinsic and BLT dsRNA-stimulated ATP hydrolysis in AncD1_VERT_ (Figure 5G). We created AncD1_VERT.7_, a construct with partial rescue of basal and BLT dsRNA-stimulated ATP hydrolysis with efficiency an order of magnitude lower than AncD1_DEUT_ and AncD1D2. Michaelis-Menten analysis indicated that the limiting factor in our rescue construct was low affinity for ATP, measured by the K_M_ value. This indicates that AncD1_VERT_ does not hydrolyze ATP, despite the conservation of the catalytic “DECH” box, because several motifs distant from the ATPase catalytic site are responsible for loss of ATP hydrolysis capability. In AncD1_VERT_ and hsDcr, these residues likely lock the helicase domain in the open conformation, preventing the formation of the ATP hydrolysis pocket in the interface between Hel1 and Hel2 (Figure 4-figure supplement 2, Figure 5). In summary, loss of ATPase function in vertebrate Dicer and consequently hsDcr, is caused by a set of mutations that restrict formation of the ATPase pocket as well as dsRNA binding and the conformational changes it would ordinarily trigger. Further engineering of AncD1_VERT_ is required to create a version of vertebrate Dicer that hydrolyses ATP more efficiently, and couples this hydrolysis to improved viral siRNA production in the context of the full-length enzyme.

In our favorite model for Dicer evolution (Figure 6), the full-length ancestral animal Dicer was capable of binding dsRNAs at two sites: the platform/PAZ domain for pre-miRNA, and HEL-DUF for long endogenous or viral dsRNA^1^. This Dicer’s helicase domain was probably more promiscuous for different dsRNA termini and possessed the ability to translocate on dsRNA. After duplication, arthropod Dicer-2 evolved to a one-site mechanism where the HEL-DUF became the primary site of dsRNA recognition, as shown in previous work where the platform/PAZ domain in arthropod Dicer-2 was observed to lose affinity for dsRNA^1^. dsRNA and ATP binding to dmDcr2’s helicase domain drives conformational changes in the entire enzyme and leads to translocation of dsRNA to the RNaseIII domain for cleavage^27,32^. On the other hand, Dicer-1 underwent a series of evolutionary changes culminating with vertebrate Dicer’s helicase domain losing affinity for both dsRNA and ATP as RLR helicases co-opted its ancestral antiviral function. Instead, vertebrate Dicer works with a one-site mechanism where the platform/PAZ domain is the predominant binding site for all dsRNAs. Further studies exploring how Dicer-1 enzymes from other invertebrates, like mollusks and echinoderms, process dsRNAs will provide a clearer picture of how different Dicer domains contribute to different RNAi pathways.

**Figure 6.**
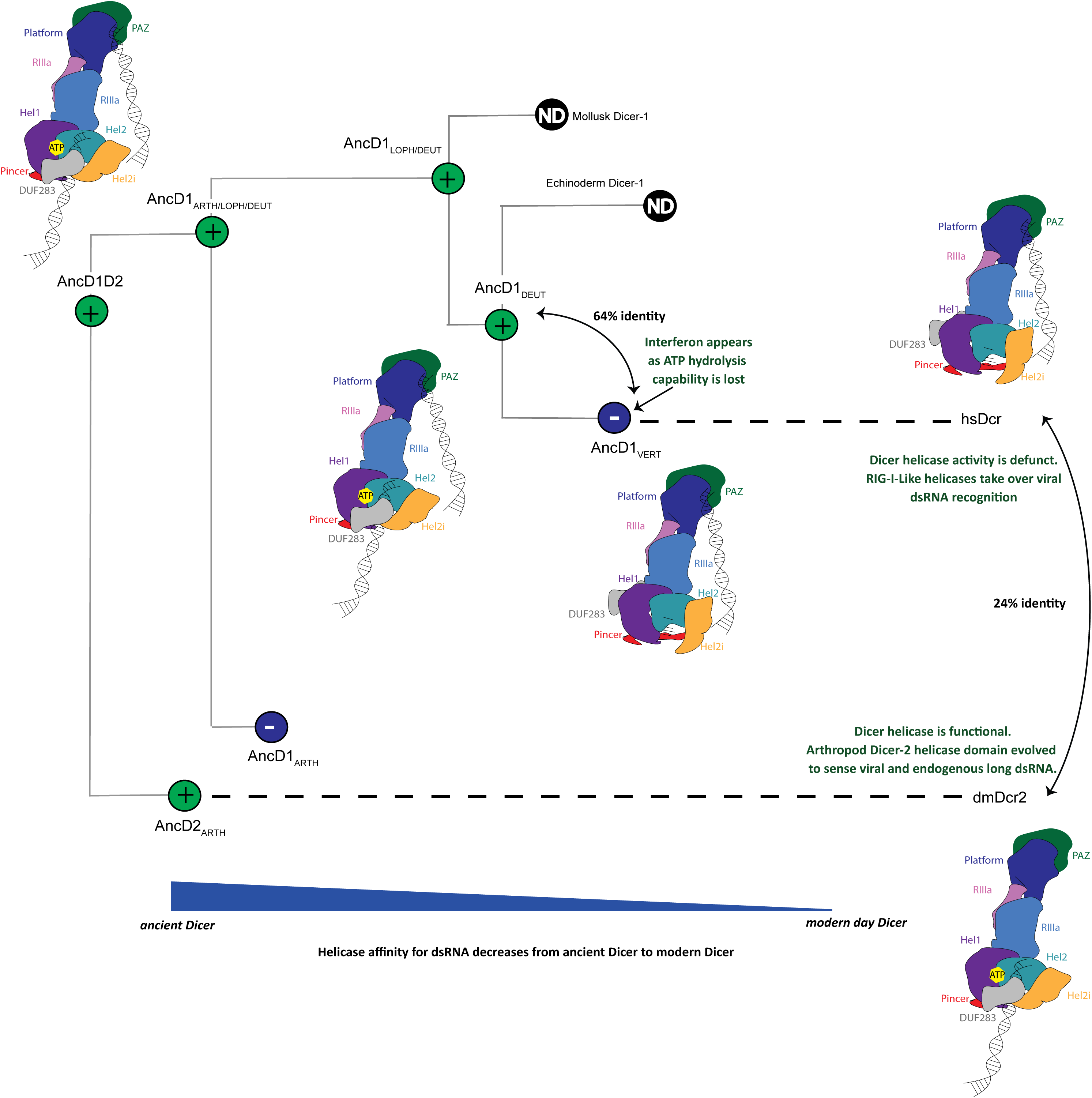
Model of metazoan Dicer evolution showing transition from a 2-site dsRNA binding in ancestral Dicer to a 1-site dsRNA binding state in extant vertebrate and arthropod Dicers. Early animals possessed one promiscuous Dicer enzyme capable of using both platform/PAZ and helicase domains for dsRNA recognition. After gene duplication, arthropod Dicer-2’s helicase domain becomes specialized for viral and endogenous long dsRNA processing and becomes the primary site of dsRNA binding. Deuterostome Dicer-1 may have retained 2-site dsRNA recognition, but at the onset of vertebrate evolution, Dicer-1 loses helicase function and exclusively uses the platform/PAZ domain to recognize dsRNA.

## Materials and Methods

### Key Resources Table

| Reagent type or resource | Designation | Source | Identifier |
| --- | --- | --- | --- |
| Expression plasmid | pFastBac-OSF | Thermo Fisher, modified in-house |  |
| Suspension cells | Spodoptera frugiperda (Sf9) cells | Expression Systems | Cat# 94-001S |
| Expression media | ESF 921 Insect Cell Media | Expression Systems | Cat# 96-001-01 |
| Bacteria cells | DH10Bac competent cells | Thermo Fisher Scientific | 10361012 |
| Gp64-PE antibody and control baculovirus | Baculovirus Titering Kit | Expression Systems | Cat# 97-101 |
| Transfection reagent | Cellfectin II | Thermo Fisher Scientific | Cat# 10362100 |

### Phylogenetics and ancestral protein reconstruction

Annotated Dicer protein sequences were retrieved from the NCBI database using taxa from each of the main animal phyla as queries for the BLAST algorithm^73^. Representative protein sequences from each metazoan phylum were used as search templates to retrieve a wide range of Dicer orthologs and paralogs with Fungus Dicers used as the outgroup to root the animal clade. Protein sequences were clustered using CD-HIT with an identity threshold of 95%, and representative sequences aligned initially with MAFFT^74,75^. Initial multiple sequence alignment (MSA) was used to assess and visualize Helicase domain and DUF283 boundaries as defined by the Conserved Domain Database^76^. All downstream analysis was performed on the Helicase and DUF283 referred to as HEL-DUF. Large gaps were manually deleted from the initial HEL-DUF alignment and PRANK was used to generate a new alignment^77^. Manual curation was carried out to remove species-specific indels and exclude sequences missing conserved parts of the helicase domain or DUF283. Model selection was performed on the resulting MSA using ProtTest 3.4.2, producing JTT+G+F as the best fit evolutionary model using the Akaike Information Criteria (AIC)^78^.

RAXML-NG v 1.0.1 was used to infer the maximum likelihood phylogeny using the best fit evolutionary model, with 8 rate categories in a gamma distribution to model among-site rate variation^79^. Transfer Bootstrap was used to calculate statistical support for the ancestral nodes with Fungal Dicers used as the outgroup for rooting the tree. Ancestral state reconstruction in RAXML-NG using the maximum likelihood tree and the JTT model^79,80^. Because an especially gappy alignment was produced as a result of using PRANK which models every unique insertion as a separate evolutionary event^77,81,82^, the input protein MSA was converted to a presence-absence alignment to model the indels in the alignment, and this matrix was used to perform ancestral protein reconstruction with the maximum likelihood phylogeny and the BINCAT model in RAXML-NG^1,83^. Overlapping protein and binary sequence ancestral reconstructions allowed the identification of spurious indels in ancestral sequences by eliminating low frequency insertions that were missed during manual curation.

### Cloning and protein expression

DNA sequences coding for select ancestral protein sequences were codon-optimized for expression in *Spodoptera frugiperda* (Sf9) insect cells using Integrated DNA Technologies’ (IDT) codon optimization tool. cDNA sequences were synthesized by IDT and subcloned into a modified pFastBac plasmid containing 2X-Strep Flag tag. Plasmids were transformed into Dh10Bac *E. coli* cells to generate bacmids, which were transfected into Sf9 cells to produce baculovirus vectors for protein expression^43^. Baculovirus titer was quantified using flow cytometry^84^. Ancestral HEL-DUF constructs were purified using Strep-Tactin Affinity chromatography, Heparin chromatography or Ion Exchange chromatography, and Size Exclusion chromatography. All ancestral constructs eluted as monomers except ANCD1_VERT_, which eluted as a mixture of monomers and dimers. Purified constructs were identified using mass spectrometry at UC Davis Proteomics Core Facility.

### dsRNA preparation

42 nt single-stranded RNAs were chemically synthesized by University of Utah DNA/RNA Synthesis Core or IDT. Equimolar amounts of single-stranded RNAs were annealed in annealing buffer (50 mM TRIS pH 8.0, 20 mM KCl) by placing the reaction on a heat block (95°C) and slow cooling ≥ 2 hours^43^. dsRNAs were gel purified after 8% polyacrylamide native PAGE and quantified using a Nanodrop.

### RNA sequences

42-nt sense RNA:

5’-GGGAAGCUCAGAAUAUUGCACAAGUAGAGCUUCUCGAUCCCC-3’

42-nt antisense BLUNT RNA:

5’-GGGGAUCGAGAAGCUCUACUUGUGCAAUAUUCUGAGCUUCCC-3’

5’-GGAUCGAGAAGCUCUACUUGUGCAAUAUUCUGAGCUUCCCGG-3’

### ATP hydrolysis assays

Reactions were performed at 37°C in 65μL reaction mixtures containing cleavage assay buffer (25mM TRIS pH 8.0, 100mM KCl, 10mM MgCl_2_, 1mM TCEP) for the times indicated, with 200nM ancestral protein and 400nM 42 BLT or 3’ovr dsRNA, in the presence of 100μM ATP-MgOAc with [α-^32^P] ATP (3000 Ci/mmol, 100nM) spiked in to monitor hydrolysis. Protein was preincubated at 37°C for 3 min prior to the addition to reaction mix. Reactions were started by the addition of protein to reaction mix containing ATP and dsRNA. 2μL of reaction were removed at indicated times, quenched by addition of 2μL of 500mM EDTA, spotted (3μL) onto 20 x 20 cm PEI-cellulose plates (Cel 300 PEI/UV 254 TLC Plates 20×20, Machery-Nagel, Ref 801063), and chromatographed with 0.75M KH_2_P0_4_ (adjusted to pH 3.3 with H_3_PO_4_) until solvent front reached the top of the plate. Plates were dried, visualized on a PhosphorImager screen (Molecular Dynamics) and quantified using ImageQuant software.

Quantification of ATP hydrolysis assays for Table 1 was done by fitting the data into a two-phase exponent, with the first phase modelled as a linear reaction between time 0 and 2.5 minutes. The first phase is considered to be a transient zero order reaction where the rate constant k is equal to the velocity of the reaction, which is the slope of ADP produced/ATP consumed (y) per minute (t). In reality, this rapid first phase probably ended before 2.5 minutes but we are limited by the nature of manually mixed assays, as opposed to stopped flow assays where mixing and signal collection can be done on the timescale of seconds.

y = kt + intercept

Data for the second phase were fit to the pseudo-first order equation y = y_o_ + A x (1-e^-kt^); where y = product formed (ADP in µM); A = amplitude of the rate curve, y_o_ = baseline, k = pseudo-first-order rate constant = k_obs_; t = time. Data points are mean ± SD (n≥3).

For the Michaelis-Menten ATP hydrolysis assays, varying amounts of ATP-MgOAc with [α-^32^P] ATP (3000 Ci/mmol, 50nM) was incubated with the indicated protein concentrations and the velocity of the steady-state reaction was calculated using a linear regression:

ADP produced (µM) = velocity (µM/min) x time (min)

The velocity recorded for each starting ATP concentration was fit into the Michaelis-Menten equation:

Velocity = V_max_ * X/(K_M_ + X), where V_max_ is the maximum enzyme velocity (µM/min), X is the ATP concentration (mM) and K_M_ is the Michaelis-Menten constant.

The turnover number, k_cat_, was calculated by dividing V_max_ by the total enzyme concentration. GraphPad Prism version 9 was used for curve-fitting analysis.

### Gel shift mobility assays

Gel mobility shift assays were performed with 20pM 42-basepair BLT or 3’ovr dsRNA, with sense strand labeled with ^32^P at the 5’ terminus. Labeled dsRNA was incubated and allowed to reach equilibrium (30 min, 4^◦^C) with HEL-DUF construct in the presence and absence of 5mM ATP-MgOAc_2_, in binding buffer (25 mM TRIS pH 8.0, 100 mM KCl, 10 mM MgCl_2,_ 10% (vol/vol) glycerol, 1 mM TCEP); final reaction volume, 20 µL. Ancestral HEL-DUF protein was serially diluted in binding buffer before addition to binding reaction. Reactions were stopped by loading directly onto a 5% polyacrylamide (19:1 acrylamide/bisacrylamide) native gel running at 200V at 4^◦^C, in 0.5X Tris/Borate/EDTA running buffer. The gel was pre-run (30 min) before loading samples. Gels were electrophoresed (2 hr) to resolve HEL-DUF-bound dsRNA from free dsRNA, dried (80^◦^C, 1 hr) and exposed overnight to a Molecular Dynamics Storage Phosphor Screen. Radioactivity signal was visualized on a Typhoon PhosphorImager (GE Healthcare LifeSciences) in the linear dynamic range of the instrument and quantified using ImageQuant version 8 software.

Radioactivity in gels corresponding to dsRNA_total_ and dsRNA_free_ was quantified to determine the fraction bound. Fraction bound = 1 – (dsRNA_free_/dsRNA_total_). All dsRNA that migrated through the gel more slowly than dsRNA_free_ was considered as bound. To determine K_d_ values, binding isotherms were fit using the Hill formalism, where fraction bound = 1/(1 + (K_d_ /[P])); K_d_ = equilibrium dissociation constant, n = Hill coefficient, [P] = protein concentration^85^. GraphPad Prism version 9 was used for curve-fitting analysis.

## Acknowledgments

We thank Alesia McKeown, Nels Elde, Paul Sigala, Peter Shen, Demian Cazalla, Tyler Starr and Thomas Koch, as well as members of the Bass Lab for helpful discussions and feedback. We thank Claudia Consalvo for help with assay design and assistance with flow cytometry. We thank James Marvin for supervision of flow cytometry procedure, and Ryan Andrews for help with figure design, and Michelle Salemi (UC Davis Proteomics Core) for assistance with protein mass spectrometry. AA was supported by the 3i Initiative Graduate Fellowship at the University of Utah. DNA synthesis and flow cytometry was performed at the University of Utah Core Facilities. This work was supported by funding to BLB from the National Institute of General Medical Sciences (R35GM141262) and the National Cancer Institute of the National Institutes of Health (R01CA260414).

---

**Figure 1-figure supplement 1.**
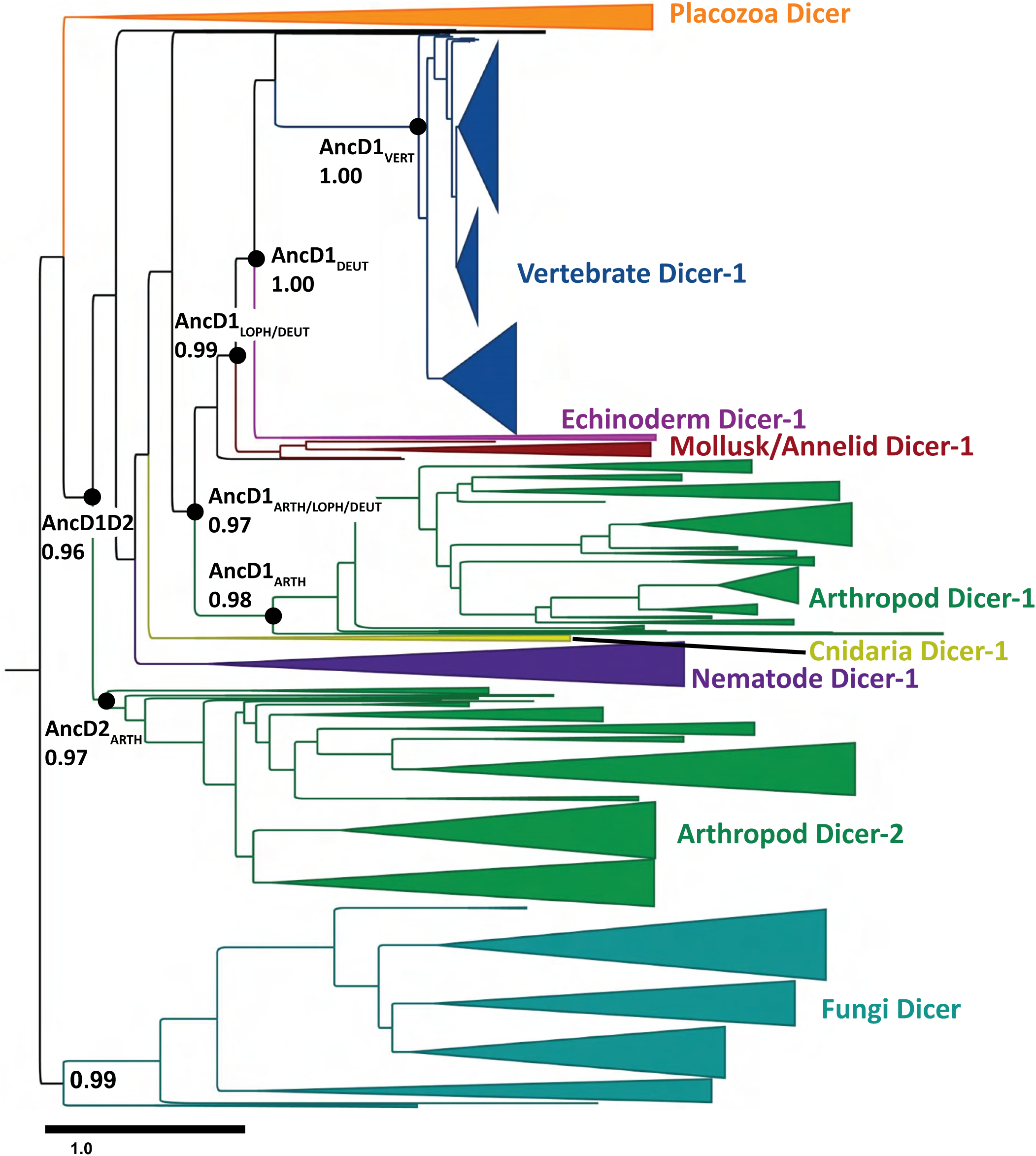
Maximum likelihood phylogenetic tree constructed from metazoan Dicer helicase domain and DUF283. Dicer HEL-DUF phylogenetic tree visualized and annotated with FigTree. Resurrected ancestral nodes are indicated by black circles, with transfer bootstrap values indicated. Width of cartoon triangle base represents number of species. Scale bar represents total amino acid substitutions divided by number of amino acid sites i.e., amino acid substitutions per site.

**Figure 1-figure supplement 2.**
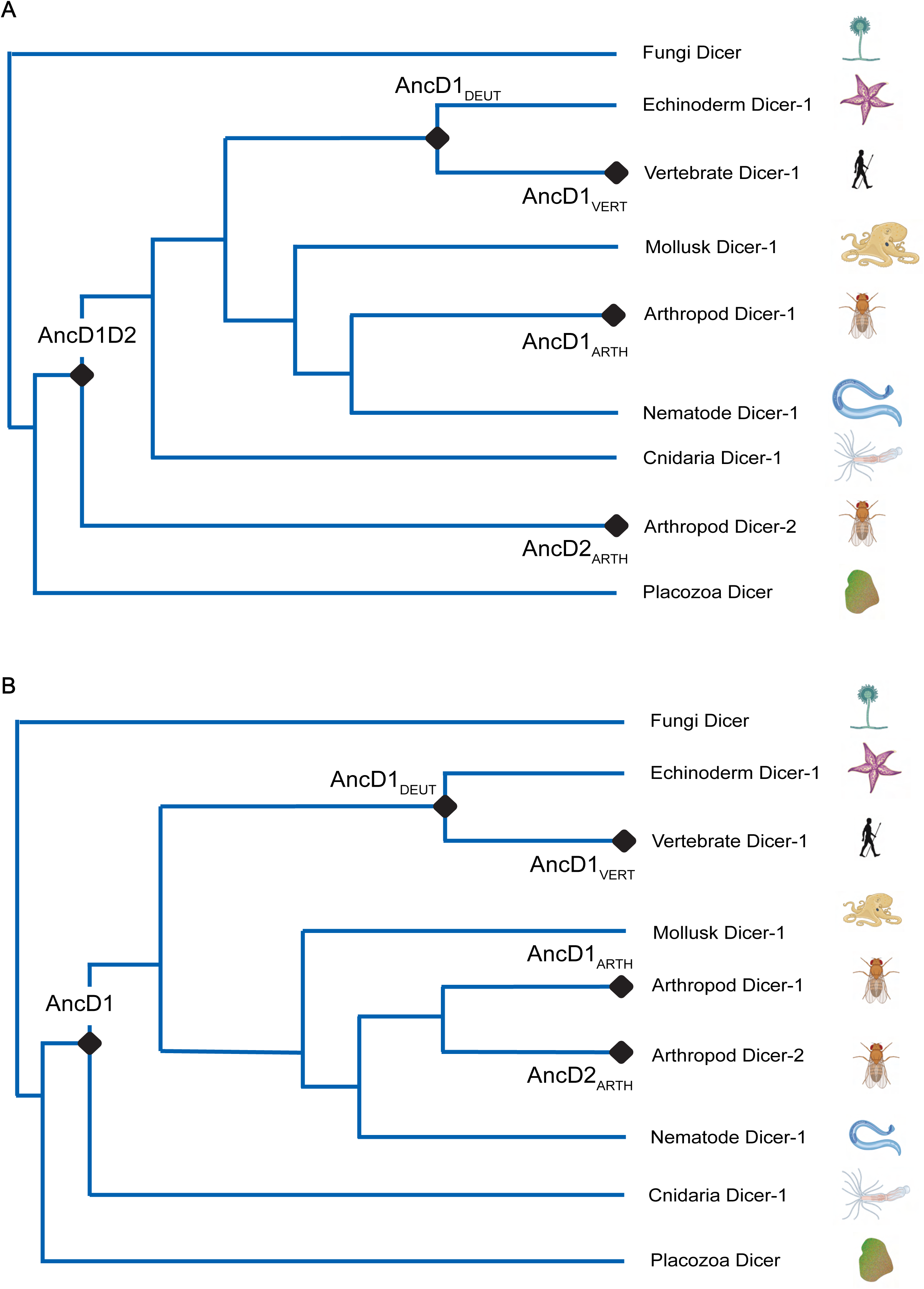
Alternative reconstructions of phylogenetic tree depicting Dicer HEL-DUF evolution. **A.** Summarized phylogenetic tree showing species-accurate relationships among metazoan phyla. Gene duplication occurs early in animal evolution. **B.** Summarized phylogenetic tree showing species-accurate relationships among metazoan phyla. Gene duplication is constrained to being arthropod-specific.

**Figure 1-figure supplement 3.**
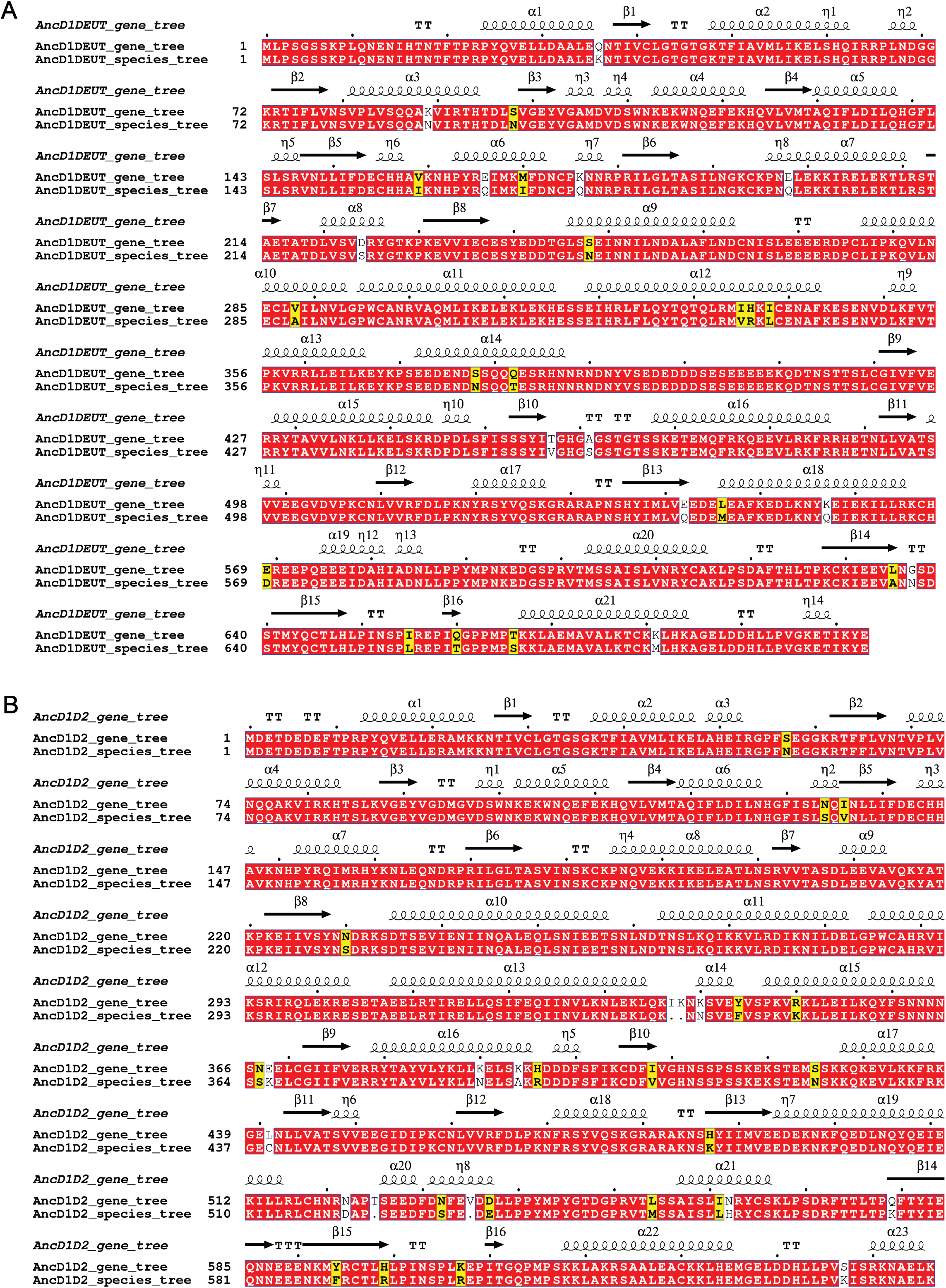
Constraining the phylogenetic tree to species-accurate relationships does not significantly impact ancestral protein reconstruction. **A.** Multiple sequence alignment illustrated with ESPript, depicting amino acid sequences for reconstructed AncD1_DEUT_ node using either the gene tree or the species tree^86^. Red, identity; yellow, similarity; unshaded, no similarity. Secondary structures for AncD1_DEUT_ determined with RosettaFold are shown above aligned sequences. TT represents beta turns. **B.** Multiple sequence alignment illustrated with ESPript, depicting amino acid sequences for the reconstructed AncD1D2 node using either the gene tree or the species tree^86^. Red, identity; yellow, similarity; unshaded, no similarity. Secondary structures for AncD1D2 determined with RosettaFold are shown above aligned sequences. TT represents beta turns

**Figure 1-figure supplement 4.**
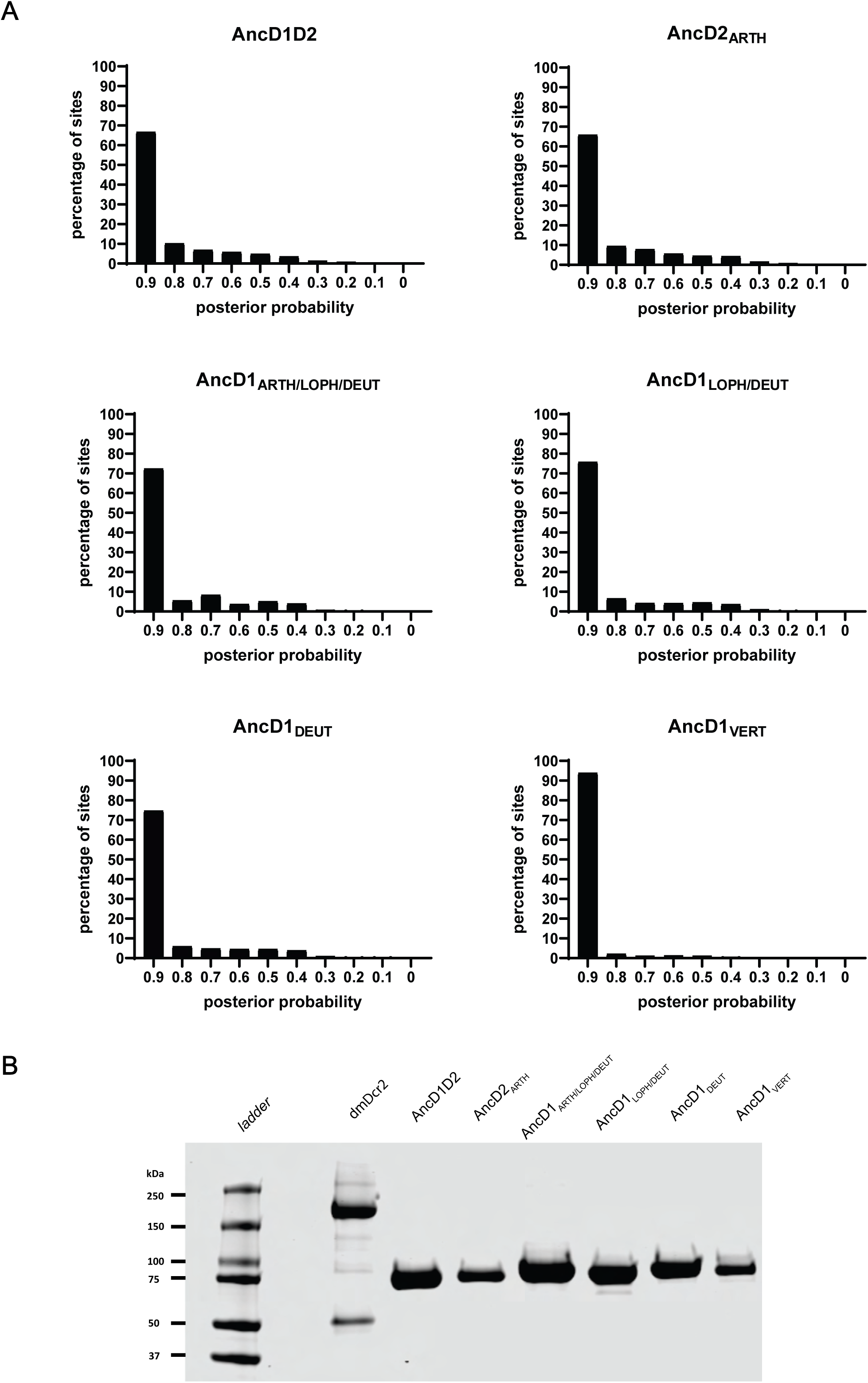
Reconstructed HEL-DUF constructs are predicted with high confidence and expressed recombinantly. **A.** Reconstructed HEL-DUFs at nodes of interest are predicted with posterior probabilities for each amino acid. Posterior probabilities for each amino acid are plotted and binned by 0.1. AncD1_VERT_ is predicted with the highest confidence. **B.** Coomassie-stained SDS-PAGE showing recombinantly expressed and purified full length *Drosophila melanogaster* Dicer-2 and ancestral HEL-DUFs.

**Figure 4-figure supplement 1.**
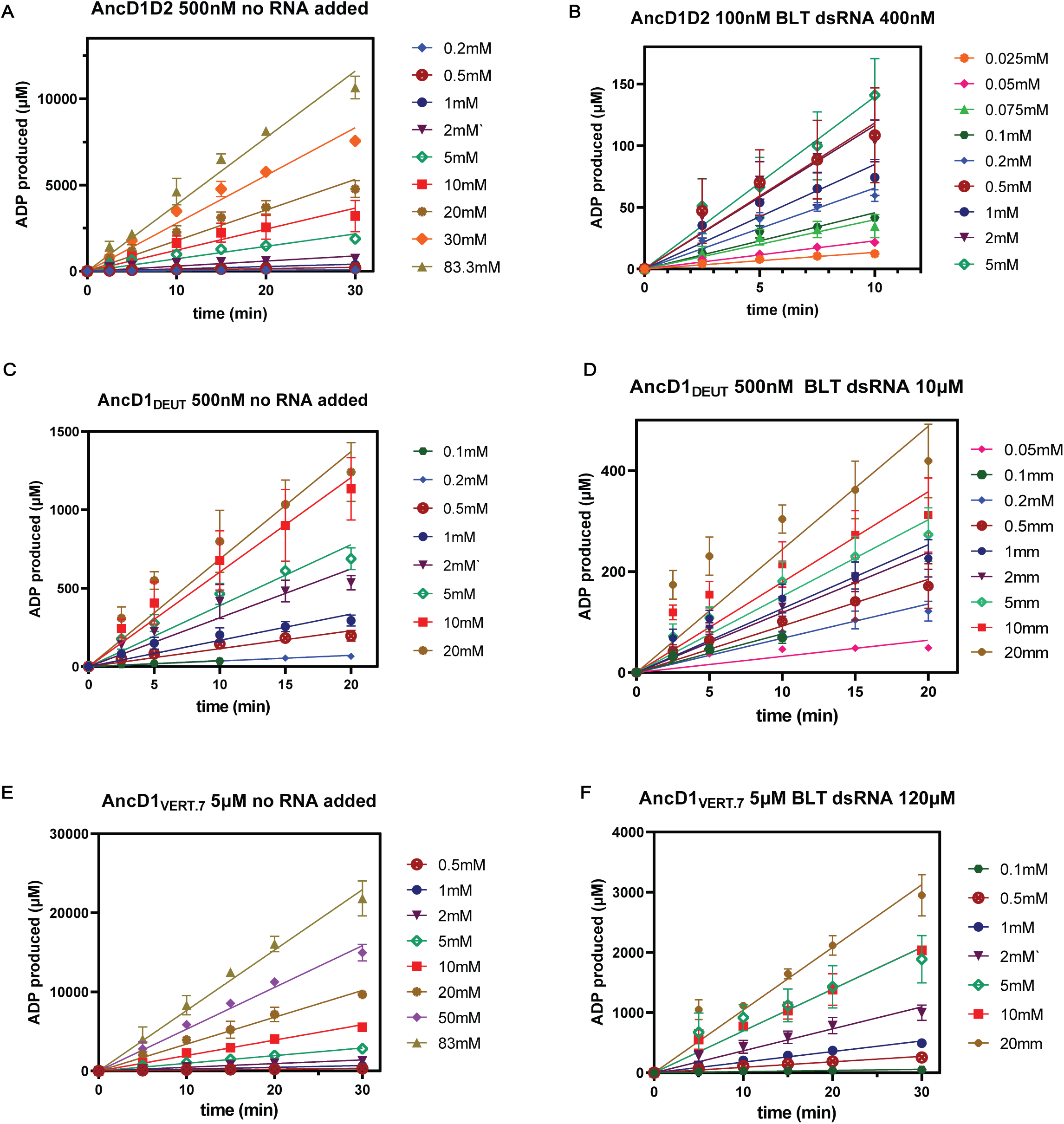
Plots of ADP production over time for ancestral HEL-DUF constructs. **A.** Basal ATP hydrolysis by 500nM AncD1D2, measured by linear ADP production over time for indicated ATP concentrations. Velocity of each reaction is the slope of the line. **B.** dsRNA-stimulated ATP hydrolysis by 100nM AncD1D2 and 400nM BLT dsRNA, measured by linear ADP production over time for indicated ATP concentrations. Velocity of each reaction is the slope of the line. **C.** Basal ATP hydrolysis by 500nM AncD1_DEUT_, measured by linear ADP production over time for indicated ATP concentrations. Velocity of each reaction is the slope of the line. **D.** dsRNA-stimulated ATP hydrolysis by 500nM AncD1_DEUT_ and 10µM BLT dsRNA, measured by linear ADP production over time for indicated ATP concentrations. Velocity of each reaction is the slope of the line. **E.** Basal ATP hydrolysis by 5µuM AncD1_VERT.7_, measured by linear ADP production over time for indicated ATP concentrations. Velocity of each reaction is the slope of the line. **F.** dsRNA-stimulated ATP hydrolysis by 5µM AncD1_VERT.7_ and 120µM BLT dsRNA, measured by linear ADP production over time for indicated ATP concentrations. Velocity of each reaction is the slope of the line. Data points, mean ± SD (n≥3).

**Figure 4-figure supplement 2.**
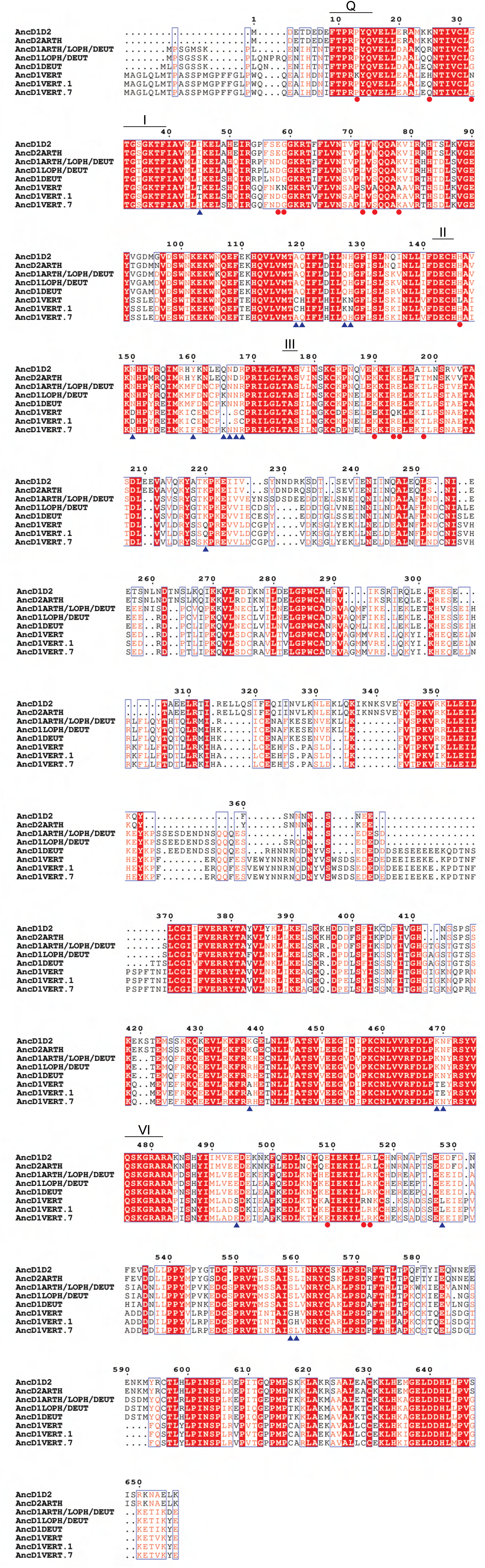
Multiple sequence alignment of ancestral HEL-DUF constructs and AncD1_VERT_ rescue constructs. Multiple sequence alignment for ancestral HEL-DUF constructs and vertebrate HEL-DUF rescue constructs, carried out with PRANK, and illustrated with ESPript. Red shading/white text indicates identity, no shading/red text indicates similarity, black text indicates no conservation. Columns with black and red text have at least 70% conservation, represented by red text, while black text indicates the non-conserved or variant amino acids. Amino acid substitutions in both rescue constructs are indicated by red circles below the column, while amino acid changes specific to AncD1_VERT_._7_ are indicated by blue triangles. Residues numbered using shortest ancestral HEL-DUF amino acid sequence. Motif Q is numbered 9-16, motif I numbered 31-38, motif II numbered 142-145, motif III numbered 175-177, motif VI numbered 476-482.

**Figure 4-figure supplement 3.**
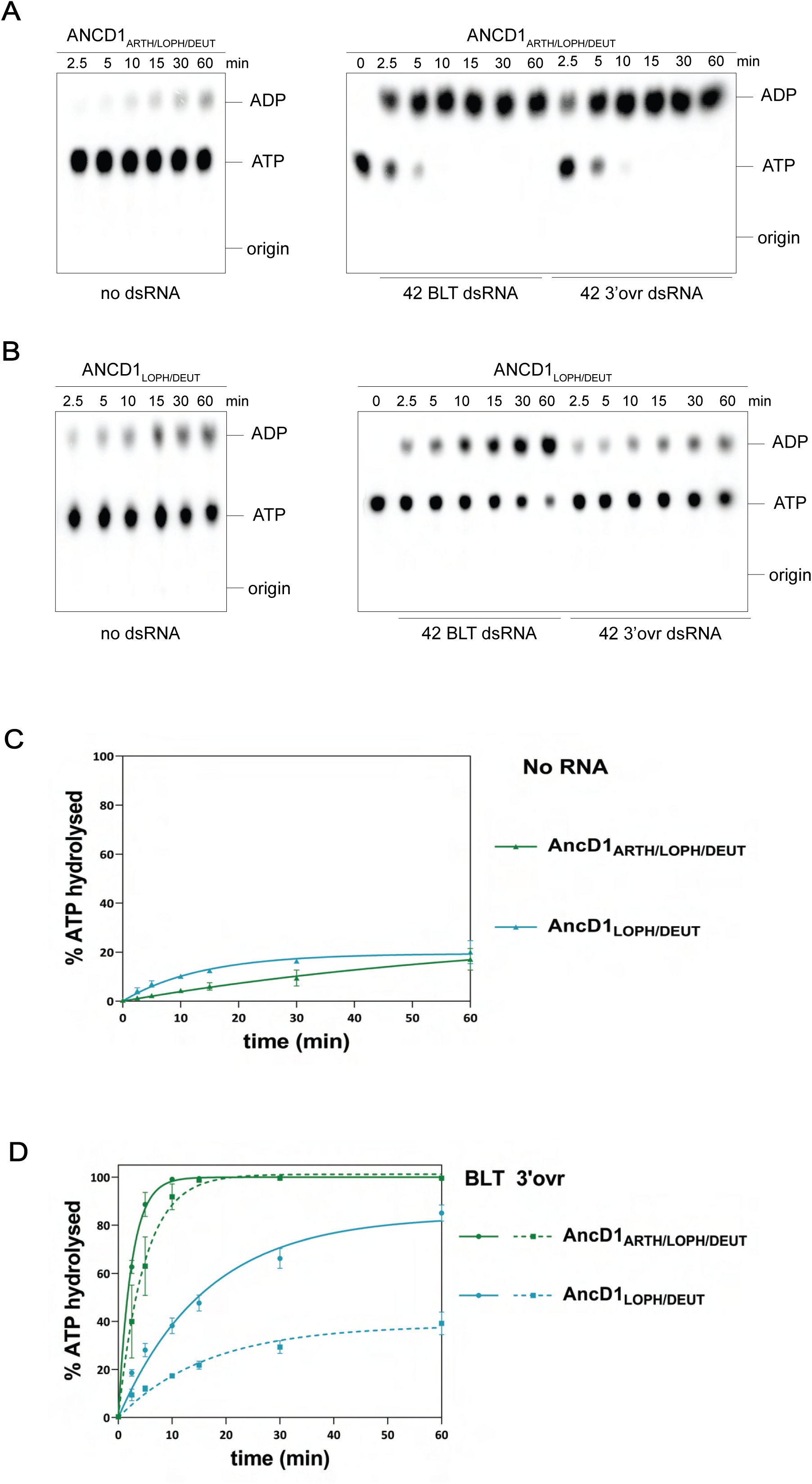
ATP hydrolysis of ancestral HEL-DUF proteins reconstructed from incongruent nodes. A-B. PhosphorImages of representative TLC plates showing hydrolysis of 100µM ATP (spiked with α-^32^P-ATP) by 200nM AncD1_ARTH/LOPH/DEUT_ (A) or AncD1_LOPH/DEUT_ (B) in absence of dsRNA (left) or in the presence of 400nM 42 base-pair dsRNA with BLT or 3’ 2-nucleotide overhang (right). **C-D.** Graph shows quantification of ATP hydrolysis assays in (A-B) performed with select ancestral HEL-DUF enzymes in the absence (C) or presence (D) of dsRNA. Data for “NO RNA” reactions were fit to the pseudo-first order equation y = y_o_ + A x (1-e^-kt^); where y = product formed (ADP in µM); A = amplitude of the rate curve, y_o_ = baseline (∼0), k = pseudo-first-order rate constant = k_obs_; t = time. Reactions with RNA were fit in two phases, first a linear phase for data below the first timepoint at 2.5 minutes, then a pseudo-first order exponential equation for remaining data. Data points are mean ± SD (n≥3).

**Figure 4-figure supplement 4.**
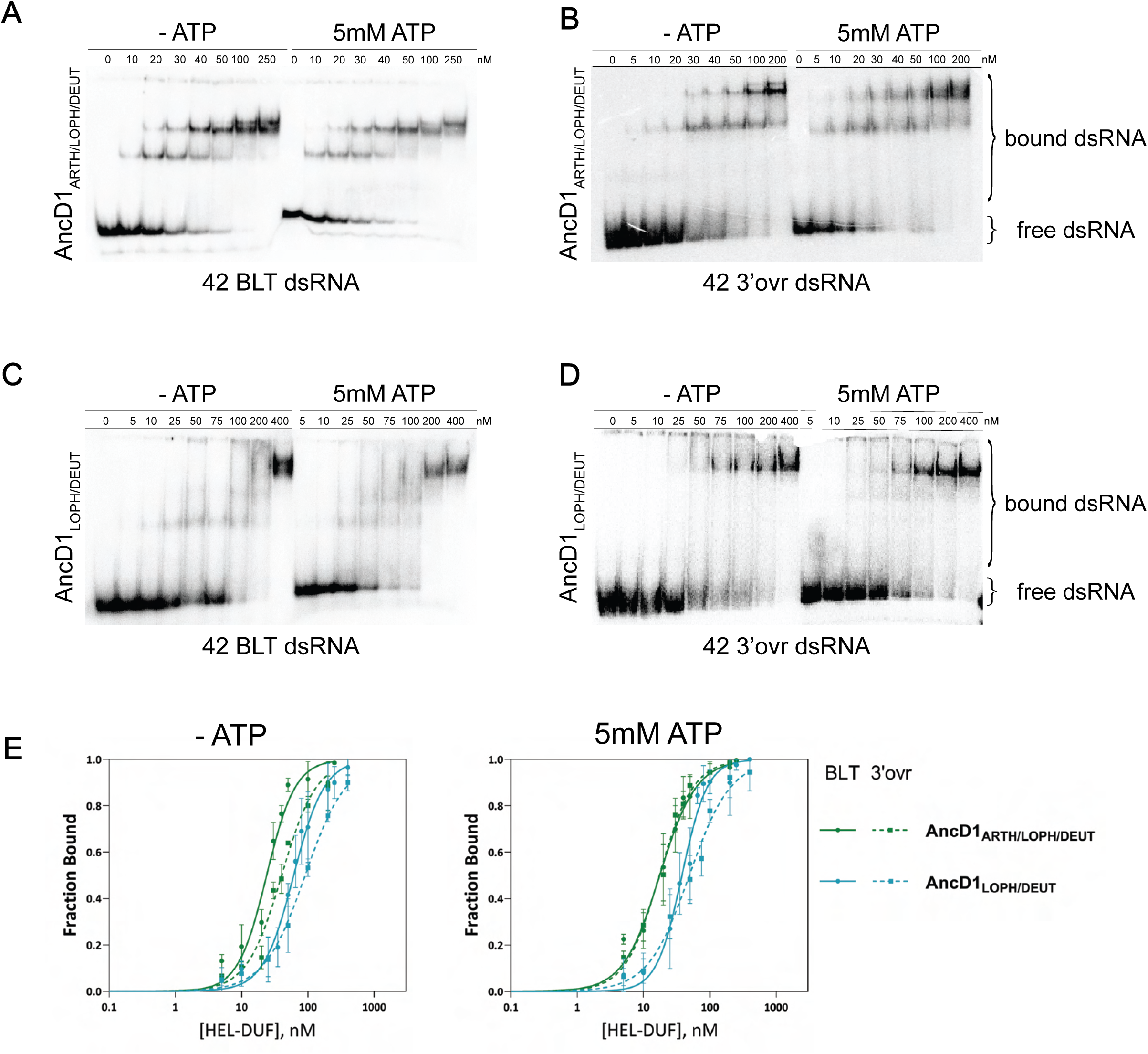
Affinity of AncD1_ARTH/LOPH/DEUT_ and AncD1_LOPH/DEUT_ for binding BLT and 3’ovr dsRNA in the absence and presence of ATP. A-D. Representative PhosphorImages showing gel mobility shift assays using select ancestral HEL-DUF constructs as indicated, and 42 base-pair BLT or 3’ovr dsRNA in the absence or presence of 5mM ATP. **E.** Radioactivity in PhosphorImages as in A-D was quantified to generate binding isotherms for ancestral HEL-DUF proteins. Fraction bound was determined using radioactivity for dsRNA_free_ and dsRNA_bound_. Data were fit to calculate dissociation constant, K_d_, using the Hill formalism, where fraction bound = 1/(1 + (K_d_ /[P])). Data points, mean ± SD (n≥3).

**Figure 1-figure supplement 4 – Source Data 1:** Original digital image of SDS-PAGE gel used in B.

**Figure 2 – Source Data 1:** Raw digital images of Thin Layer Chromatography plate used in 2A.

**Figure 2 – Source Data 2**: Raw digital image of Thin Layer Chromatography plate used in 2A.

**Figure 2 – Source Data 3:** Raw digital image of Thin Layer Chromatography plate used in 2B.

**Figure 2 – Source Data 4**: Raw digital image of Thin Layer Chromatography plate used in 2B.

**Figure 2 – Source Data 5**: Raw digital image of Thin Layer Chromatography plate used in 2C.

**Figure 2 – Source Data 6**: Raw digital image of Thin Layer Chromatography plate used in 2C.

**Figure 2 – Source Data 7**: Raw digital image of Thin Layer Chromatography plate used in 2D.

**Figure 2 – Source Data 8**: Raw digital image of Thin Layer Chromatography plate used in 2D.

**Figure 3 – Source Data 1**: Raw digital image of Gel Shift phosphoimager plate used in 3B.

**Figure 3 – Source Data 2**: Raw digital image of Gel Shift phosphoimager plate used in 3C.

**Figure 3 – Source Data 3**: Raw digital image of Gel Shift phosphoimager plate used in 3C.

**Figure 3 – Source Data 4**: Raw digital image of Gel Shift phosphoimager plate used in 3D.

**Figure 3 – Source Data 5**: Raw digital image of Gel Shift phosphoimager plate used in 3E.

**Figure 3 – Source Data 6**: Raw digital image of Gel Shift phosphoimager plate used in 3F.

**Figure 3 – Source Data 7**: Raw digital image of Gel Shift phosphoimager plate used in 3G.

**Figure 4-figure supplement 3 – Source Data 1:** Raw digital image of Thin Layer Chromatography plate used in Figure 4-figure supplement 3A, left panel.

**Figure 4-figure supplement 3 – Source Data 2:** Raw digital image of Thin Layer Chromatography plate used in Figure 4-figure supplement 3A, right panel.

**Figure 4-figure supplement 3 – Source Data 3:** Raw digital image of Thin Layer Chromatography plate used in Figure 4-figure supplement 3B.

**Figure 4-figure supplement 3 – Source Data 4:** Raw digital image of Thin Layer Chromatography plate used in Figure 4-figure supplement 3B.

**Figure 4-figure supplement 4 – Source Data 1:** Raw digital image of Gel Shift phosphoimager plate used in Figure 4-figure supplement 4A.

**Figure 4-figure supplement 4 – Source Data 2:** Raw digital image of Gel Shift phosphoimager plate used in Figure 4-figure supplement 4B.

**Figure 4-figure supplement 4 – Source Data 3:** Raw digital image of Gel Shift phosphoimager plate used in Figure 4-figure supplement 4C.

**Figure 4-figure supplement 4 – Source Data 4:** Raw digital image of Gel Shift phosphoimager plate used in Figure 4-figure supplement 4D.

## Notes

### Competing Interest Statement

The authors have declared no competing interest.

